# Transcriptome profiling of a multiuse model species *Lymnaea stagnalis* (Gastropoda) for ecoimmunological research

**DOI:** 10.1101/2020.09.23.308643

**Authors:** Otto Seppälä, Jean-Claude Walser, Teo Cereghetti, Katri Seppälä, Tiina Salo, Coen M. Adema

## Abstract

Host immune function can contribute to numerous ecological/evolutionary processes. Ecoimmunological studies, however, typically use one/few phenotypic immune assays and thus do not consider the complexity of the immune system. Therefore, “omics” resources that allow quantifying immune activity across multiple pathways are needed for ecoimmunological models. We applied short-read based RNAseq (Illumina NextSeq 500, PE-81) to characterise transcriptome profiles of a multipurpose model species *Lymnaea stagnalis* (Gastropoda). We used a genetically diverse snail stock and exposed individuals to immune elicitors (injury, bacterial/trematode pathogens) and changes in environmental conditions that can alter immune activity (temperature, food availability). Immune defence factors identified in the *de novo* assembly indicated uniform aspects of molluscan immunity: pathogen-recognition receptors (PRR) and lectins activate Toll-like receptor (TLR) pathway and cytokines that regulate cellular and humoral defences. However, also apparent differences to other taxa were detected (i.e., modest numbers of antimicrobial peptides and fibrinogen related proteins). Identified factors also indicate that several of them might contribute to the phenotypic immune assays used on this species. Experimental treatments revealed factors from non-self recognition (lectins) and signalling (TLR pathway, cytokines) to effectors [e.g., antibacterial proteins, phenoloxidase (PO) enzymes] whose gene expression depended on immune activations and environmental conditions, as well as components of snail physiology/metabolism that may drive these effects. Interestingly, gene expression of many factors (e.g., PRR, lectins, cytokines, PO enzymes, antibacterial proteins) showed high among-individual variation. Such factors are important to include in ecoimmunological research because they may explain among-individual differences in parasite resistance and fitness in natural populations.

## 1. Introduction

Several fields of ecology and evolution increasingly recognise host immune function as an essential contributor to biological processes (see Schmid-Hempel 2011). For instance, the immune system plays critical roles in life-history evolution (Zuk and Stoehr 2002; Nunn et al. 2009), sexual selection (Hamilton and Zuk 1982; Rantala et al. 2000), and responses/adaptations of organisms to environmental change (Siva-Jothy and Thompson 2002; Wilson et al. 2002; Leicht et al. 2017). This recognition has given rise to an interdisciplinary field of ecological immunology (or ecoimmunology; see Demas and Nelson 2012). Ecoimmunological studies, especially in invertebrates, typically measure the end products of one or few immunological cascades that are controlled by several genes (e.g., Cotter et al. 2004; Schwarzenbach et al. 2005). Thus, the field relies on quantitative genetic theory, initially motivated by the assumed simplicity of innate-type invertebrate immune systems with non-specific recognition and killing mechanisms. Over the recent decades, however, comparative immunology with aid from genomics, has shown invertebrate immune systems to be complex and diversified across the phyla (e.g., Loker et al. 2004; Ghosh et al. 2011; Deleury et al. 2012; Dheilly et al. 2014; Doublet et al. 2017). In fruit flies, for instance, specific immune pathways respond towards Gram-positive and Gram-negative bacteria, as well as fungi (Hoffmann and Reichhart 2002; Royet et al. 2005; Hetru and Hoffmann 2009).

The realised complexities of invertebrate immune systems provide new challenges and opportunities for studies focusing on ecological and evolutionary questions on immune function. This is because typical ecoimmunological studies that focus on one/few immunological mechanisms are incapable of describing the multivariate “immune phenotypes” (*sensu* Pedersen and Babayan 2011) of organisms. That, however, would be important because each immunological pathway can respond differently to various selective agents (e.g., parasite/pathogen species) and may be traded-off with different physiological, life-history, and other immune traits (Cotter et al. 2004; Hanelt et al. 2008; Adema et al. 2010; Deleury et al. 2012; Zhang et al. 2015). Additionally, the expression of immune traits, as well as the associated trade-offs, may depend on environmental conditions such as resource availability and temperature (e.g., Feder et al. 1997; Lee et al. 2006; Leicht et al. 2017), which could be due to stress-related responses or altered metabolism. Therefore, a comprehensive characterisation of different immunological and physiological mechanisms, as feasible by genomics-type approaches, in ecologically relevant experiments utilising invertebrates is essential. This expansion has been done successfully in some insects such as bumblebee (e.g., Barribeau et al. 2014; Brunner et al. 2014) and red flour beetle (Ferro et al. 2017; Greenwood et al. 2017). Considering the extensive diversity of invertebrate phyla, development and utilisation of such genomic data also in other taxonomic groups would further expand the potential of ecoimmunology.

Mollusca is the second-largest animal phylum after Arthropoda. In this phylum, Gastropoda represents the largest taxonomic class with 40,000 to 150,000 living species (Lindberg et al. 2004). Gastropods inhabit aquatic and terrestrial habitats, and incur disease from viruses and bacteria (e.g., Moore et al. 2002; Prince 2003; Travers et al. 2008) but also from specialist flatworm parasites called digenetic trematodes (Cribb et al. 2003; Olson et al. 2003). Trematodes have complex life cycles that typically involve a gastropod as an intermediate host. In snails, trematodes reproduce asexually to produce transmission stages that infect the next host in the life cycle (another intermediate host or a definitive host). Most of the attention on molecular immunology in gastropods has focused on *Biomphalaria glabrata* (Planorbidae, Hygrophila, Panpulmonata), a tropical freshwater snail that transmits the human blood fluke, *Schistosoma mansoni* (Bayne 2009; Coustau et al. 2015; Adema et al. 2017). Attention to other species has been comparatively limited. However, several gastropods, including pond snails of the family Lymnaeidae (Hygrophila, Panpulmonata), transmit medically and veterinary-relevant parasites in temperate regions. These include liver fluke (*Fasciola hepatica*), fish eye flukes (*Diplostomum* spp.), and bird schistosomes that cause swimmer’s itch (e.g., Caron et al. 2014; Horák et al. 2015; Selbach et al. 2015). Nevertheless, diversity and function of circulating defence cells, haemocytes, have been investigated in the great pond snail, *Lymnaea stagnalis* (van der Knaap et al. 1981; 1993; Adema et al. 1994; Horák and Deme 1998; Lacchini et al. 2006; Wright et al. 2006; Walker et al. 2010; Skála et al. 2020), and a draft genome of this species is available (Davison et al. 2016).

Here, we applied short-read based RNAseq to characterise transcriptome profiles of *L. stagnalis* exposed to various immune elicitors (injury, bacterial and trematode pathogens) and environmental changes (temperature, food availability) using a genetically diverse laboratory population of snails. *Lymnaea stagnalis* is a model organism in multiple biological disciplines (reviewed in Fodor et al. 2020), including ecological immunology (e.g., Seppälä and Jokela 2010, 2011; Langeloh et al. 2017; Leicht et al. 2017; Salo et al. 2017; 2019). Earlier ecoimmunological work, however, employs only a few phenotypic immune assays. Therefore, our transcriptome annotation analyses focused on identifying a broad range of candidate immune genes to develop approaches that can track various immune mechanisms for future ecoimmunological research on this species, as well as to widen the comparative immunology of molluscs. Furthermore, we evaluated variation in expression of the identified candidate genes to detect immunological mechanisms that respond to particular immune elicitors/environmental conditions and/or that show high among-individual variation. We propose that the use of such genes as targets would allow a comprehensive examination of polymorphic immune function in future ecoimmunological studies.

## 2. Materials and methods

### 2.1 Study animals and experimental design

This study employed adult *L. stagnalis* snails (*N* = 48, shell length 24.6–30.5 mm) from an F_2_ generation of a laboratory stock population that was generated by interbreeding snails that originated from seven different natural populations in Switzerland (see Langeloh et al. 2017). Interbreeding was done to increase genetic variation among the experimental snails relative to natural populations that may show low genetic diversity (Kopp et al. 2012). Higher genetic diversity among experimental snails increased the potential to identify immune genes that show variation in their expression levels among individuals depending, for example, on their genotype and allelic variation.

Before the experiment, experimental snails were placed individually in perforated 0.2 L plastic cups and distributed in four water baths each containing 40 L of aged tap water at 20°C, with twelve snails per water bath. Snails were fed with fresh lettuce *ad libitum*, and half of the water in each water bath was changed every second day. This set-up allowed water changes with minimal disturbance to the snails. Snails were acclimated to these conditions for four (treatments including an environmental change) to six days (treatments including an immune challenge) before exposure to experimental manipulations (see the next two paragraphs for the description of the treatments). The acclimation period differed between treatment types to allow simultaneous sampling of snail tissues for RNA extractions (i.e., duration of the treatments was not the same).

In the experiment, the snails were randomly assigned to nine experimental treatments (Table 1). In seven treatments, snails’ responses to immune elicitors, including the effects of handling that were necessary for immune challenge treatments, were tested. Snails in treatment one (untreated controls; *N* = 5) were maintained as during the acclimation period. Snails in treatment two (anaesthetised controls, *N* = 5) were anaesthetised by placing them into 2% diethyl ether for two and a half minutes. These snails were used to examine the effects of snail handling when compared to untreated controls (treatment 1 above) because anaesthesia was needed in following immune challenge treatments. In treatment three (wounding; *N* = 6), each anaesthetised snail was injected with 170 μL of snail saline (see Standen 1975) in the head-foot using a syringe and needle (0.45 mm × 16 mm BD Microlane™ 3, New Jersey, USA). This treatment examined the effect of wounding in activating immune defence when compared to anaesthesia without exposure to immune elicitors (treatment 2 above). The sham injection also served as a control for the treatments involving bacterial injections (described in the next paragraph).

**Table 1.**
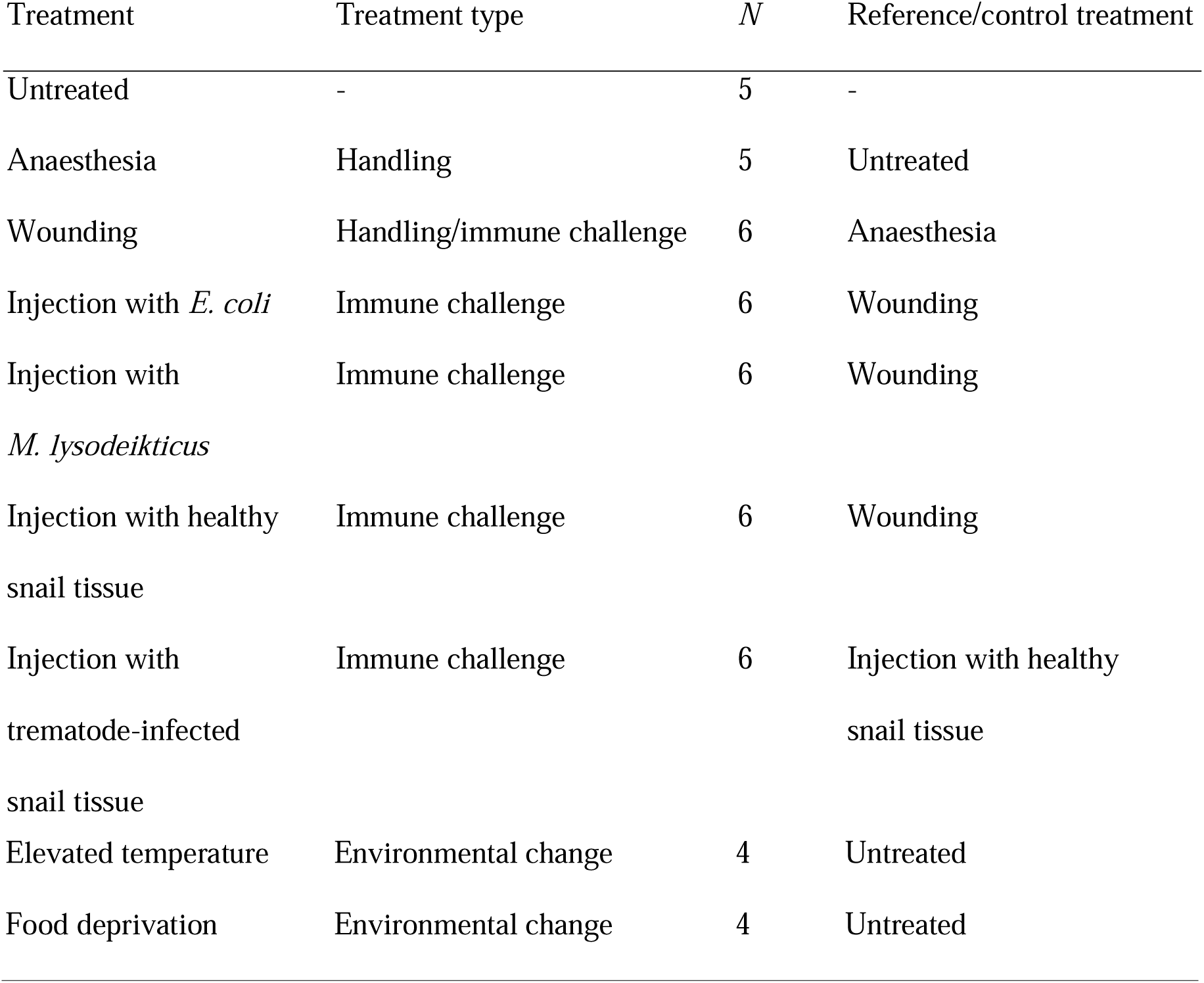
Summary of the experimental design presenting the treatments, treatment types (handling used in immune challenges, immune challenge, environmental change), the number of exposed snails, and the specific reference/control treatment used for each treatment group when examining their effects on gene expression.

In treatments four and five (bacterial injections; *N* = 6 for each), anaesthetised snails were injected as above with 0.5 mg/mL lyophilised *Escherichia coli* (Sigma-Aldrich, Steinheim, Germany; product number EC11303) or *Micrococcus lysodeikticus* (synonym *Micrococcus luteus*; Sigma-Aldrich, Steinheim, Germany; product number M3770) cells in snail saline, respectively. Both Gram-negative (*E. coli*) and Gram-positive (*M. lysodeikticus*) bacteria were used because lipopolysaccharides of Gram-negative bacteria and peptidoglycans of Gram-positive bacteria are recognised by different receptors that activate pathogen-specific immune responses (Poltorak et al. 1998; Takehana et al. 2002; Montminy et al. 2006; Hanelt et al. 2008; Deleury et al. 2012).

In treatments six and seven (snail/trematode-tissue injections; *N* = 6 for each), anaesthetised snails were injected as above with soluble extracts from healthy or trematode-infected gonads (ovotestes) dissected from *L. stagnalis* snails collected from a pond in Winterthur, Switzerland (47° 28’N, 8° 43’E). In donor snails, parasite infections were evident from the shedding of strigeid and echinostome-type cercariae. Both healthy and infected gonads were disrupted in 1.5 mL Eppendorf tubes using plastic pestles, adding 1 mL of snail saline per 100 mg of tissue. Samples were vortexed and pelleted briefly (2000 × g for 5 seconds). The supernatant was collected from three uninfected and three infected snails and the samples from each snail type were pooled for injections. The treatment using the extract from healthy gonads served as a control for injection with tissue extract from the trematode-infected snail. After injections, all experimental snails were placed back in their original cups with lettuce for six hours to allow their immune systems to respond to immune elicitors.

In addition to immune challenges, snails’ responses to environmental changes were tested in two treatments. Snails in treatment eight (elevated temperature; *N* = 4) were placed in water baths where the water temperature was raised to 25°C during several hours and maintained at an elevated level for two days (see Salo et al. 2017; 2019). After the transfer, additional snails were added to all water baths to keep the experimental snails continuously at a density of twelve snails per water bath (data were not collected from these extra snails). Snails in treatment nine (food deprivation; *N* = 4) were maintained without food for two days. Untreated snails (treatment 1 above) served as controls for these treatments. All snails survived the experimental treatments and displayed normal motility and activities during the experiment.

### 2.2 RNA extraction, library preparation and sequencing

At the end of the experimental treatments, each snail was removed from its shell, and soft body tissues were submerged in self-made RNAlater solution (see protocol at https://www.protocols.io/view/RNAlater-Recipe-c56y9d) in which they were cut into small pieces. The cut tissues were stored in 10 mL of RNAlater at −20°C. Within the next 15 days, all samples were disrupted under liquid nitrogen using mortar and pestle. Total RNA was extracted from samples of tissue grinds (71-102 mg per snail) using TRIzol reagent (Invitrogen, Carlsbad, CA, USA) according to the manufacturer’s instructions except for conducting RNA wash with 75% EtOH three times. Residual genomic DNA was removed from samples using Turbo DNA-free kit (Ambion, Austin, TX, USA) according to the manufacturer’s instructions. RNA quantity was measured using Qubit 2.0 Fluorometer (Invitrogen, Carlsbad, CA, USA), and sample quality and purity were verified using 2100 Bioanalyzer (Agilent Technologies, Palo Alto, CA, USA; RNA Nano chips) and NanoDrop ND-1000 spectrophotometer (NanoDrop Technologies Inc., Wilmington, DE, USA), respectively. Complementary DNA (cDNA) libraries were constructed for each snail individual using TruSeq Stranded mRNA Sample Preparation Kit (Illumina, San Diego, CA, USA) according to the manufacturer’s instructions. All libraries were sequenced on Illumina NextSeq 500 platform using paired-end reads with 81 nt read length at the Genomics Facility Basel.

### 2.3 RNA-seq data handling and transcriptome assembly

Raw Illumina reads from all libraries were subjected to adaptor trimming and quality filtering. Reads with Illumina TruSeq adaptors were trimmed using Cutadapt v1.5 (min. overlap 30 nt and max. allowed error rate 0.05). Reads were then quality filtered using PRINSEQ-lite v0.20.4 [min. read length: 50 nt, GC range: 10-90%, mean Q ≥ 5, trim base left and right with Q < 10, no ambiguous nucleotides allowed, trim poly A/T tail (min. 5) from both sequences, and remove duplicate reads]. Finally, Phix sequences with BenBank genome reference NC_001422.1, as well as SSU and LSU rRNA sequences with SILVA (Release 111) reference, were removed using BBDuk v2015.08.21. To generate a *L. stagnalis de novo* reference transcriptome, the cleaned reads from nine libraries (one randomly chosen library per experimental treatment) were combined and assembled with trinity v2.0.6 (parameters: min contig length: 100, min glue: 4, min kmer cov: 4, group pairs distance: 300, path reinforcement distance: 85, normalise reads, normalise max read cov: 50, and normalise by read set). Likely contaminant transcripts from other organisms were removed by mapping all transcripts on the *L. stagnalis* reference genome (BenBank accession number GCA_900036025.1) and removing those that lacked a significant similarity hit in the genome and that were not previously characterised from *L. stagnalis*. Completeness of the produced reference transcriptome was assessed by examining the detection of core BUSCO genes (Simão et al. 2015; Waterhouse et al. 2018) from metazoans (978 genes).

### 2.4 Annotation of immune-, stress- and metabolism-related factors

Amino acid sequences of previously characterised proteins/genes relevant for immune function, stress responses and metabolism primarily from molluscs (especially gastropods; e.g., Gorbushin and Borisova 2015; Adema et al. 2017), but also from other organisms, were collected from GenBank (Table S1) and used in BLAST (v2.6.0+) similarity searches to identify orthologs in the *L. stagnalis* reference transcriptome. Components of the immune system queried included non-self recognition through pathogen-recognition receptors (PRRs; all abbreviations are explained at the end of the article), Toll-like receptor (TLR) pathway, cytokines, antibacterial peptides and proteins, production of reactive oxygen species (ROS), phenoloxidase (PO)/melanisation-type defence, and apoptosis. Components of stress responses included cell protection and survival, oxidative stress and antioxidant enzymes. Targeted components linked to metabolism included regulatory proteins, proteins related to the transport of elements and compounds as well as a selection of receptors.

From the statistically best Blast hits (E-value ≤ 0.05), transcripts that were at least 60% of the length of the shortest reference sequence and showed minimum 40% similarity were used to identify open reading frames (ORFs, EMBOSS v6.6.0.0) that were between a start and a stop codon. ORFs that were at least 60% of the length of the shortest reference sequence (absolute minimum length: 30 amino acids) were examined for the presence of a signal peptide (SignalP v4.1) and the domain structure (SMART, PFAM, NCBI). For identification of fibrinogen-related proteins (FREPs), sequences that were computationally identified to contain fibrinogen-like (FBG) domains were manually inspected for upstream immunoglobulin superfamily (IgSF) domains because the diffuse sequence motifs of invertebrate IgSF domains frequently challenge automated detection.

### 2.5 Gene expression analysis

To examine variation in the expression levels of different transcripts among experimental snails, sequencing reads from each library were first indexed to transcripts in the reference transcriptome using Kallisto v.0.44.0 with 100 bootstraps (Bray et al. 2016). Sleuth was then used for downstream processing (Pimentel et al. 2017). The transcriptome-wide expression profiles in different libraries were visualised using their principal component (PC) scores obtained from a principal component analysis (PCA, the first 5 PCs were used) using the internal normalization in Sleuth. Very high variation in the expression profiles among snail individuals even within experimental treatments (see the Results section) negated the use of formal statistical tests for variation in the expression of individual transcripts. Therefore, the expression levels of individual transcripts that represented the annotated factors were visualised using heatmaps that presented all experimental snails. Signal strength was calculated in units of transcripts per million (TPM). This measure calculates the proportion of counts per base for each transcript in the whole data set (multiplied by million).

## 3. Results

### 3.1 Sequencing and assembly

Illumina sequencing of the libraries yielded a total of 1.08 billion paired-end 81 nt long reads (PE-81). The raw reads representing a total of 175.2 Gbp sequence data were deposited in the NCBI Sequence Read Archive (accession number PRJNA664475). The *de novo* assembly of the reads from nine libraries (one per each experimental treatment) yielded 264,746 transcripts with a N50 of 1,589 nt, mean length 551 nt, and median length of 188 nt (146.0 Mb in total). Using TransDecoder (v3.0.1), 68,473 ORFs longer than 100 aa were identified. The removal of likely contaminant sequences that lacked a significant similarity hit with *L. stagnalis* genome (GenBank accession number GCA_900036025.1) and that were not previously characterised from this species, yielded a final reference transcriptome (139.1 Mb) consisting of 226,116 contigs with a mean transcript length of 615 nt (assembly available in https://doi.org/10.5281/zenodo.4044169). BUSCO analysis indicated high completeness of the reference transcriptome based on detection of 98.8% (complete plus partial) of the set of metazoan universal single-copy orthologs (complete: 96.3% [duplicated: 28.2%], fragmented: 2.5%, missing: 1.2%, *N* = 978).

### 3.2 Sequences supporting species identification

The obtained reference transcriptome included sequences that represent parts of the snail rDNA cassette, including the complete ITS1 and ITS2 regions (GenBank accession numbers MT864603 and MT864602, respectively). *Lymnaea stagnalis* was confirmed as the species identity of the experimental snails by the highest sequence similarities (BLASTN) of these sequences with GenBank entries from this species. Similarly, snails exposed to tissue extracts from trematode-infected gonads provided LSU-derived sequences with the highest similarity with Clinostomidae and Plagiorchiidae families of trematodes (GenBank accession numbers MT872505 and MT872506, respectively), which confirms the morphological identification of cercaria released by the donor snails.

### 3.3 Transcriptome annotation

The mining of the *L. stagnalis* reference transcriptome yielded numerous factors that contribute to immune responses from non-self recognition to the elimination of pathogens (Fig. 1, Table S2). For accommodation of PRRs, *L. stagnalis* expressed seven variants (i.e., transcripts with unique ORFs) of long-form (no short forms) peptidoglycan recognition proteins (PGRPs), four variants of Gram-negative bacteria binding-proteins (GNBPs) and a repertoire of lectins. Representatives of the detected lectin families included two FREPs of the VIgLs (variable immunoglobulin and lectin-domain-containing molecules), galectins comprising of either one, two or four carbohydrate recognition domains, multiple Chi-lectins, as well as L- and M-type lectins. The latter two families may, however, function mostly in housekeeping roles.

**Figure 1.**
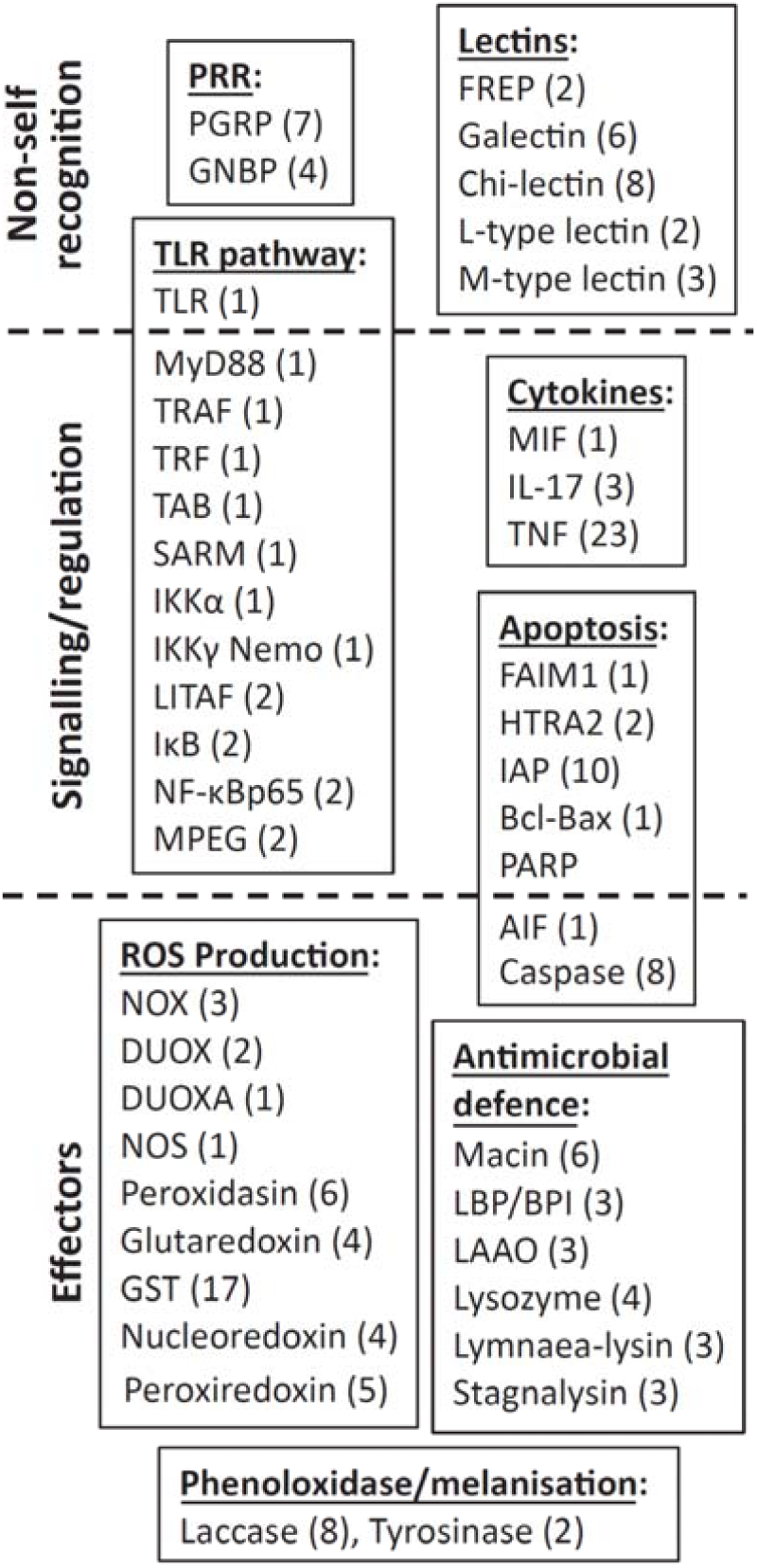
Summary of the identified immune defence factors in *Lymnaea stagnalis* reference transcriptome organized across different immunological mechanisms/pathways and steps of the immune response (i.e., non-self recognition, signalling/regulation, effectors). Numbers in brackets show how many transcripts with unique open reading frames (ORFs) were detected from those factors for which determining the completeness of ORFs was possible.

Along with one TLR sequence encoding the canonical architecture of tandemly arranged leucine-rich repeats (LRRs), a transmembrane region and a C-terminal Toll-interleukin receptor (TIR) domain, the associated signalling NFκB pathway for immune activation was represented by multiple components such as adaptor proteins for intracellular signalling downstream of the receptor (MyD88, TRAF), transcription factor (Rel protein) NF-κB, and various regulators (e.g., SARM, IκB). Additionally, three categories of cytokines for intercellular communication (including immune responses and inflammation) were detected; one variant of macrophage migration inhibitory factor (MIF), three variants of interleukin 17A (IL-17) and 23 variants of tumor necrosis factor (TNF).

The search for antimicrobial defences yielded a single family of macin antimicrobial peptides (AMPs, 6 variants), as well as several families of antimicrobial proteins, with an abundant representation of lipopolysaccharide-binding protein/bactericidal permeability-increasing proteins (LBP/BPI, 3 variants), L-amino acid oxidases (LAAOs, 3 variants), lysozymes (4 variants), and transcripts encoding for cytolytic β pore-forming toxins. These latter sequences were designated Lymnaea-lysins (3 variants) and stagnalysins (3 variants), in analogy to the orthologous Biomphalysins and glabralysins described from *B. glabrata* (Galinier et al. 2013; Lassalle et al. 2020). Central components from a gene network facilitating the production and management of reactive oxygen species for oxidative killing mechanisms were detected, including the membrane-bound enzyme complex NADPH-oxidase (NOX, with Cytochrome B 245 as the main component, 3 variants) that produces superoxide anions, as well as dual oxidases (DUOX, 2 variants) that catalyse the synthesis of anion superoxide and hydrogen peroxide. Moreover, dual oxidase maturation factor (DUOXA, 1 variant) that activates DUOX was found, as well as additional proteins involved in the biosynthesis of other oxidative compounds. Furthermore, eight variants of laccase and two of tyrosinase that can contribute to PO/melanisation-type defence were recorded. The search also yielded transcripts encoding for antioxidant enzymes and proteins regulating oxidative damage. These transcripts included two types of superoxidase dismutase enzymes against ROS [SOD (2 variants) and MnSOD (1 variant)], catalase (CAT, 2 variants), glutathione peroxidase (3 variant), and glutathione reductase (1 variant).

Our analysis also revealed transcripts that encode for proteins involved in apoptosis (programmed cell death), a process that also functions in internal defence responses. Identified transcripts included FAIM1 that regulates the extrinsic signalling pathway leading to apoptosis. Representatives of the intrinsic signalling pathway included serine protease HTRA2 (2 variants) that contribute to the loss of the mitochondrial transmembrane potential, one apoptosis-inducing factor (AIF) that causes DNA fragmentation and chromatin condensation, and factors that regulate these processes [Baculovirus IAP Repeat domain-containing caspase inhibitors, Bcl-Bax, PARP]. Finally, eight variants of caspases that destroy critical cellular proteins were detected.

The search for response factors related to environmental change yielded two families of heat shock proteins that are induced by a variety of stressors: HSP70 (5 variants) and HSP90 (1 variant). Along with the HSPs, a heat shock factor (HSF, 1 variant) that regulates the expression of HSPs was identified. Additionally, our analysis yielded transcripts linked to invertebrate metabolism. These included one protein phosphatase (PP1) that controls for cellular processes linked to metabolism, gene transcription and translation, cell movement and apoptosis, and eight variants of ubiquitin, a regulatory protein that marks proteins, for instance, for degradation and recycling by proteasome and affects protein activity. Furthermore, transcripts encoding for ferritin (3 variants) that participates in iron transport and storage in invertebrates and protects organisms from iron-induced oxidative stress, and alcohol dehydrogenase (ADH, 1 variant) that catalyses the interconversion between alcohols and aldehydes or ketones were identified. Lastly, the search yielded transcripts linked to the estrogen-related receptor (ERR, 2 variants) and retinoid acid receptor (RXR, 3 variants) that regulate, for instance, oxidative metabolism, energy homeostasis, and imposex development in invertebrates.

### 3.4 Transcriptomic responses to experimental treatments

Plots of PCA scores for the transcriptome-wide expression profiles of individual snails did not reveal any grouping based on experimental treatments or outlier libraries that deviated from others (Fig. 2, Fig S1). Heatmaps for the expression levels of the annotated immune factors (see Table S2) across experimental treatments indicated some immune-elicitor specific responses. For instance, injection with lyophilised *E. coli* cells increased the gene expression of TLR, as well as of some components of the TLR signalling pathway (IκB, NF-κB), when compared to the expression levels in snails injected with physiological saline (comparison 1 in Fig. 3, Fig. S2). Additionally, exposure to *E. coli* increased the expression levels of effectors representing antibacterial defence [Lymnaea-lysins (4 out of 6 individuals); comparison 2 in Fig. 3, Fig. S3] and PO/melanisation-type reaction (laccase; comparison 3 in Fig. 3, Fig. S4). Exposure of snails to lyophilised *M. lysodeikticus* cells activated the expression of the lectin FREP (3 out of 6 individuals; comparison 4 in Fig. 3, Fig. S5), the cytokine IL-17 (comparison 5 in Fig. 3, Fig. S6), one component of the TLR signalling pathway (IκB; comparison 6 in Fig. 3, Fig. S2), and the effector laccase (comparison 3 in Fig. 3, Fig. S4). The injection of soluble extracts from trematode-infected snail tissue increased the expression level of laccase (comparison 7 in Fig. 3, Fig. S4) when compared to the snails challenged with extracts from healthy snail tissue. Also, IκB was activated in some individuals exposed to extracts of both healthy and trematode-infected snail tissue (comparison 8 in Fig. 3, Fig. S2). Additionally, wounding led to upregulation in the expression of DUOXA (Fig. S7) that contributes to ROS production. These responses conform to the notions of their importance for invertebrate immune function.

**Figure 2.**
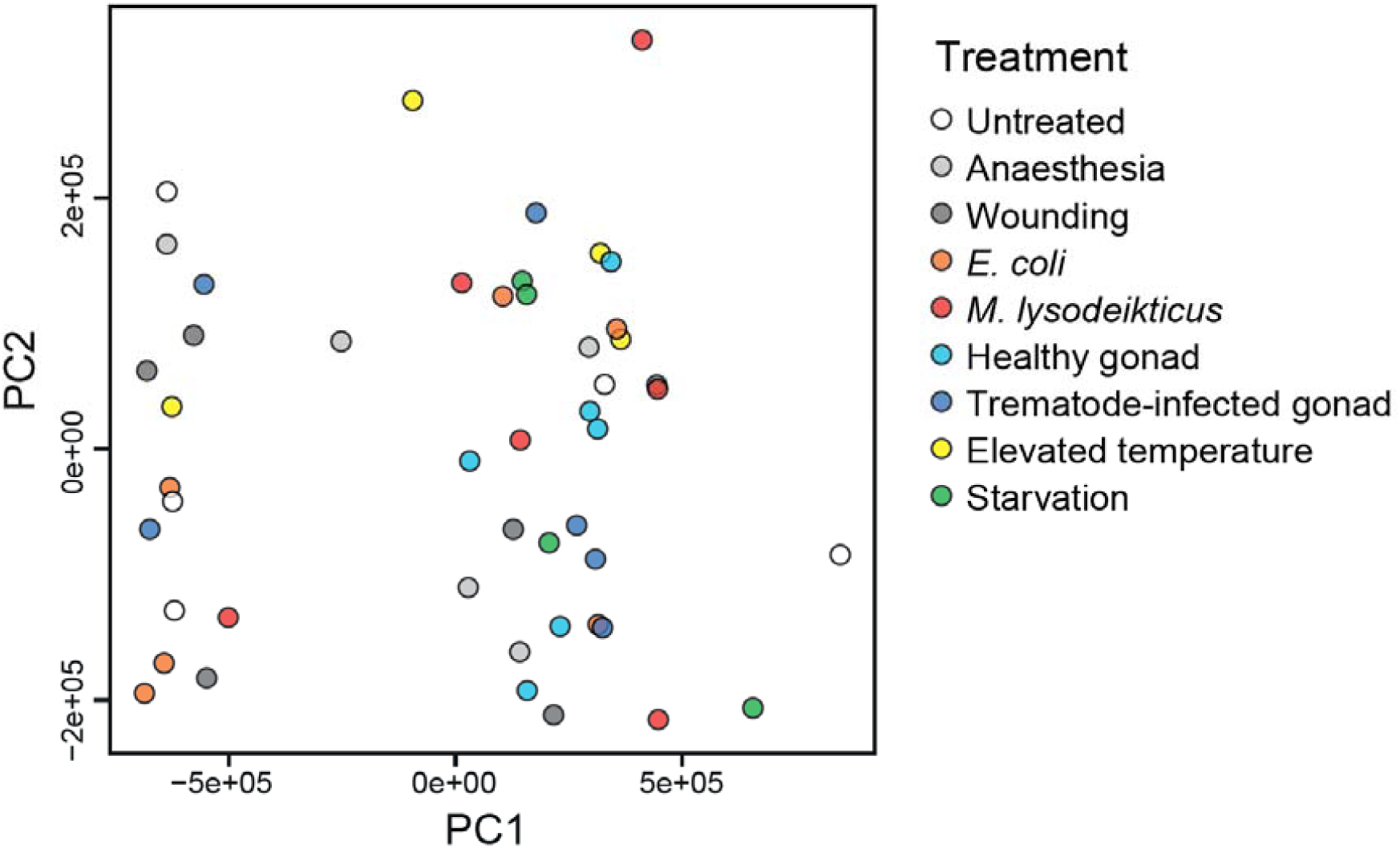
A principal component analysis (PCA) plot showing the variation in transcriptome-wide expression profiles of the experimental snails using the first two principal components (PCs) after internal normalization in Sleuth. PCA plots for the first five PCs, as well as the proportion of total variance each of them explained in the data, are presented in Fig. S1.

**Figure 3.**
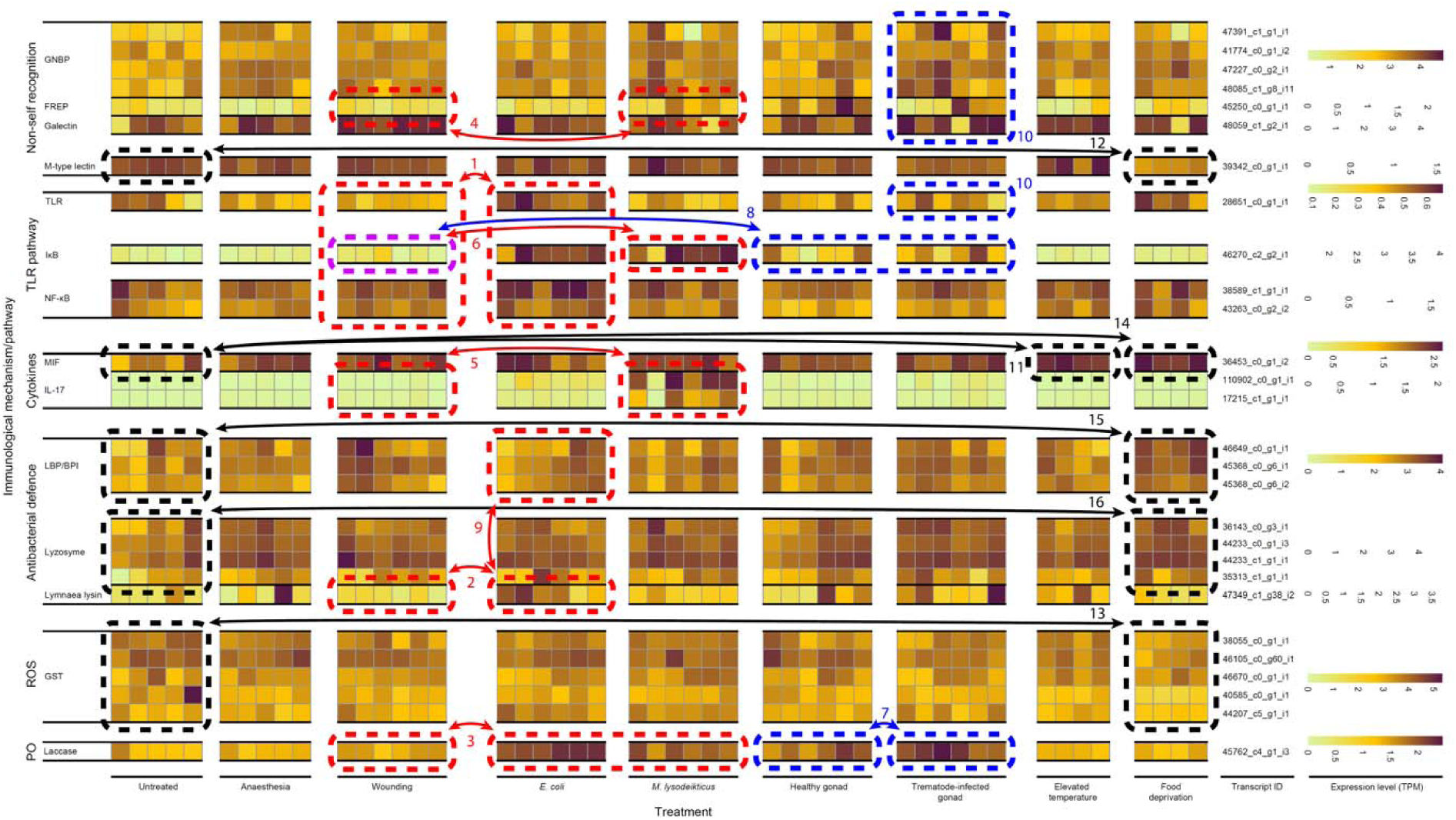
Expression levels of individual transcripts representing different immunological pathways/mechanisms [non-self recognition, TLR signalling, cytokines, antibacterial defence, production of reactive oxygen species (ROS), phenoloxidase (PO)/melanisation-type reaction] that showed deviations in certain experimental treatments from their specific controls or individuals expressed distinct transcripts within certain components of the immune system (transcripts related to each factor are clustered according to their similarity). Heatmap shows the variation for each factor among all experimental snails (each column represents one snail) using its dynamic range in units of transcripts per million (TPM). Red (injections with bacteria), blue (injections with snail/trematode tissue extracts), purple (injections with bacteria and snail/trematode tissue extracts) and black (exposures to environmental change) rectangles (dashed line), arrows and numbers refer to the specific results mentioned in the text.

Befitting the innate type immunity of invertebrates, the expression levels of other immune factors did not show systematic differences among immune activation treatments (Figs. S2-S8). However, the expression levels of many factors showed high among-individual variation within treatment groups. This was the case for PRRs GNBP (Fig. S5), galectin (Fig. S5) and TLR (Fig. S2), as well as for some components of the TLR signalling pathway (Myd88, TRAF, NF-κB p65; Fig. S2), cytokines MIF and TNF (Fig. S6), several components of ROS production (NOX, DUOXA, GST, nucleoredoxin, NOS, peroxidasin; Fig. S7), the effector tyrosinase (PO/melanisation-type reaction; Fig. S4), factors regulating apoptosis (HTRA2, AIF, Bcl-Bax, PARP; Fig. S8), and all examined antibacterial defence factors (Fig. S3). A few immune factors showed high among-individual variation that was limited to certain immune activation treatments: lectin FREP (Fig. S5), IκB (TLR pathway; Fig. S2), DUOX (ROS production; Fig. S7), laccase (PO/melanisation-type reaction; Fig. S4) and the cytokine IL-17 (Fig. S6). Interestingly, within certain components of the immune system, individual snails expressed distinct gene transcripts, which suggests different defence strategies against pathogens. For instance, snails challenged with *E. coli* showed high gene expression in two to four of the examined six antibacterial defence factors and the factors with high expression levels were different among individuals (Fig. S3). Most strikingly, individuals with the highest expression of Lymnaea lysins showed the lowest expression of LBP/BPI and vice versa (comparison 9 in Fig. 3). Similarly, snails exposed to trematode-infected snail tissue extracts showed high expression levels in only one to two of the examined non-self recognition components (GNBP, FREP, galectins, TLR), and the expressed factors were different among individuals (areas 10 in Fig. 3).

The examined environmental changes apart from immune challenge affected several components of the snail immune system. The elevated temperature increased the expression of the cytokine MIF (comparison 11 in Fig. 3, Fig. S6) when compared to untreated snails. Food deprivation, on the other hand, downregulated the expression of an M-type lectin (comparison 12 in Fig. 3, Fig. S5), and glutathione S-transferase (GST, comparison 13 in Fig. 3, Fig. S7), but led to upregulation of MIF (comparison 14 in Fig. 3, Fig. S6), LBP/BPI (comparison 15 in Fig. 3, Fig. S3) and lysozyme (comparison 16 in Fig. 3, Fig. S3) gene expression. Additionally, some of the factors that contribute to stress-responses, antioxidation and metabolism were found to be affected by the experimental treatments. First, heat shock proteins (both HSP70 and HSP90; comparison 1 in Fig. 4, Fig. S9) and glutathione reductase (comparison 1 in Fig. 4, Fig. S10) showed reduced expression levels under food deprivation. Exposure of snails to immune elicitors (injection with lyophilised *E. coli* and *M. lysodeikticus* cells, injection with healthy snail gonad) increased the expression of ADH (comparison 2 in Fig. 4, Fig. S11) when compared to the snails injected with physiological saline. Similarly to the immune defence genes, high among-individual variation was seen in some annotated factors with a potential role as antioxidant enzymes [SOD (all treatments), MnSOD (some treatments), glutathione reductase (most immune activation treatments); Fig. S10] or in snail metabolism [PP1 (most treatments), ADH (most immune activations), RXR, (most treatments); Fig. S11].

**Figure 4.**
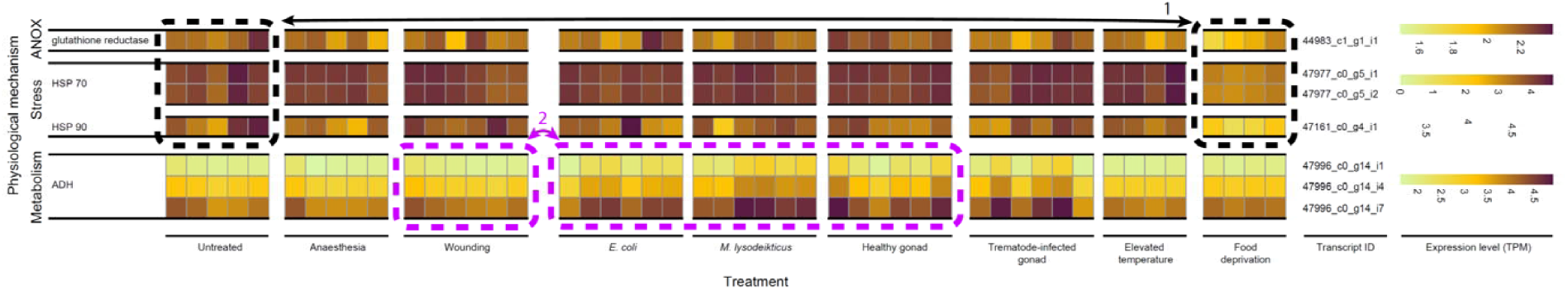
Expression levels of individual transcripts representing antioxidant enzymes (ANOX), as well as proteins related to stress responses and metabolism that showed deviations in certain experimental treatments from their specific controls (transcripts related to each factor are clustered according to their similarity). Heatmap shows the variation for each factor among all experimental snails (each column represents one snail) using the dynamic range in units of transcripts per million (TPM). Purple (immune challenge) and black (exposures to environmental change) rectangles (dashed line), arrows and numbers refer to the specific results mentioned in the text.

## 4. Discussion

This study aimed to comprehensively describe the transcriptomic responses of the freshwater snail *L. stagnalis* to immune challenges and environmental changes, and thus to produce a resource for future ecoimmunological research. Earlier work on snail ecoimmunology has relied on quantifying a narrow subset of phenotypic immune defence traits. Those traits, however, respond differently to immune elicitors (Seppälä and Leicht 2013) and environmental conditions (e.g., Seppälä and Jokela 2010; Leicht et al. 2013; Leicht et al. 2017; Salo et al. 2017), and they contribute differently to snail fitness (Langeloh et al. 2017). Additionally, contrary to the earlier expectation of simple innate-immune system in invertebrates (i.e., non-specific defence mechanisms) the recent development in comparative immunology and genomics/transcriptomics has revealed invertebrate immune systems to be highly complex, diverse, and also specific against different parasite types (e.g., Loker et al. 2004; Ghosh et al. 2011; Deleury et al. 2012; Dheilly et al. 2014; Doublet et al. 2017). Together these findings call for ecological experiments that quantify snail “immune phenotypes” across a wide range of immunological mechanisms. Only such studies may evaluate the role of snail immune function as a whole in ecological and evolutionary processes.

In this study, the samples for transcriptome sequencing were extracted from whole body tissues of individual snails to avoid any bias in the recovery of immune and stress-response factors that are expressed in specific tissues or cell types (see e.g., Baron et al. 2016; Adema et al. 2017). The reference transcriptome assembled from nine individuals exposed to different experimental treatments may not fully represent the genome content of *L. stagnalis* because mRNA-based approaches only capture genes that are actively expressed. BUSCO analysis, however, indicated representation of 98.8% of the universally shared metazoan genes in the assembly. Therefore, the reference transcriptome can be assumed to be highly comprehensive. Additionally, the use of a genetically diverse lab stock may have led to the capture of allelic variants of genes that do not all occur within single snail individuals, yet exist in natural populations where they may contribute differently to organismal fitness. This is supported by the high proportion (28.2%) of core BUSCO genes that were found with more than one copy.

Along with the aim to support future ecoimmunological research, the characterisation of immune genes of *L. stagnalis* broadens comparative immunology of aquatic pulmonate snails (Hygrophila), previously available only for a few species from the families Lymnaeaidae, Physideae and Planorbidae (Adema et al. 2017; Schultz and Adema 2017; Alba et al. 2019). The organisation of antimicrobial defences in *L. stagnalis* is in line with hygrophilid snails that all show considerable numbers of antimicrobial proteins (LBP/BPI, LAAO, lysozyme, lymnaealysin, cytolytic β pore-forming toxins), contrasted by a modest number of macin type AMP genes (5 in *L. stagnalis*). This is remarkable because of the great diversity and numbers of AMP genes and gene families that are usually recorded from other organisms, including bivalve molluscs (e.g., Mitta et al. 2000). Perhaps hygrophylid gastropods employ novel categories of AMPs that remain to be characterised, but it appears that these snails have a unique approach to dealing with invading microbes.

The recovery of *L. stagnalis* FREPs is consistent with the distribution of VIgLs as likely multimeric immune recognition factors across Gastropoda (Gorbushin 2019). The low number of FREPs in *L. stagnalis* is similar to that of the physid *Physella acuta* (Schultz et al., 2017) but differs from the planorbid *B. glabrata* that generates inter-individually unique repertoires of FREPs (Zhang et al. 2004). Furthermore, the transcriptomic responses of *L. stagnalis* to the introduction of extracts of healthy gonad tissue from other individuals (used as a control for injection with trematode-infected gonad extracts) was remarkably similar to those elicited by wounding alone. The lack of an immune response to allotypic tissue extracts suggests the absence of (acute) allograft rejection in *L. stagnalis*, just as it lacks from closely related planorbid snails (Sullivan et al. 1995; 1998). Together these findings suggest that snail immune function has evolved in a lineage-specific manner. However, our study supports the view that the main aspects of innate immunity are conserved throughout animal evolution (Buchmann 2014). For example, lectins and GNBP receptors are available to detect pathogens and signal through the TLR pathway and cytokines to activate defence responses that encompass cellular (e.g., ROS production) and humoral (antimicrobial proteins) branches of the immune system.

The obtained detailed information regarding the molecular immunology of *L. stagnalis* can now be integrated into future ecoimmunological investigations. Studies on *L. stagnalis* typically measure two phenotypic immune defence traits from snail haemolymph, namely antibacterial activity and PO-like activity (e.g., Seppälä and Jokela 2010, 2011; Leicht et al. 2013; Langeloh et al. 2017; 2017; Salo et al. 2017; 2019). Antibacterial activity is quantified as a reduction in optical density of a solution in which lyophilised *E. coli* cells are mixed with snail haemolymph (see Seppälä and Jokela 2010; Seppälä and Leicht 2013). The current study identified several different effector proteins that may contribute to this reaction. These included AMPs, LBP/BPI, LAAOs, lysozymes and cytolytic β pore-forming toxins. Therefore, the phenotypic antibacterial activity assay is likely to estimate the combined activity of these components. The other commonly studied phenotypic immune defence trait, PO-like activity, measures an increase in optical density of a solution in which PO enzymes from snail haemolymph oxidise the substrate L-dopa (see Seppälä and Jokela 2010; Seppälä and Leicht 2013). This assay may also measure the combined activity of different factors (see Le Clec’h et al. 2016). Our study revealed two categories of PO enzymes, laccase and tyrosinase, suggesting that they may be the most important contributors to the melanisation-type reaction in *L. stagnalis*. Further tests at the phenotypic level using enzyme-specific substrates can estimate their relative importance in snail immune function (see Le Clec’h et al. 2016).

Various immune activation treatments and manipulations of environmental conditions were used in this study to identify genes that contribute to snail immune responses against different pathogen types (bacteria, trematode parasites) and that are affected by environmental (ambient temperature, resource availability) variation. Visual examination of the gene expression patterns of the annotated factors indicated different responses both against Gram-negative (*E. coli*) and Gram-positive (*M. lysodeikticus*) bacteria. Exposure to *E. coli* activated TLR pathway as well as the gene expression of two types of effectors, Lymnaea-lysins (a component of antibacterial activity) and laccase (an enzyme with PO activity). Exposure to *M. lysodeikticus* led to upregulation in one FREP lectin and cytokine IL-17. Exposure to trematode-infected snail tissue extracts activated only the gene expression of laccase. The observed patterns suggest specificity against different types of pathogens by some components of *L. stagnalis* immune system and are in line with earlier findings in other invertebrates (e.g., Hoffmann and Reichhart 2002; Royet et al. 2005; Hanelt et al. 2008; Hetru and Hoffmann 2009; Deleury et al. 2012). Two of the annotated factors, however, showed broader responses to the utilised immune elicitors, thus suggesting their more general roles in immune function. First, laccase that may be a key component of PO activity in molluscs (Le Clec’h et al. 2016) responded both to *E. coli* and extracts of trematode-infected snail tissue. Second, one of the central regulators of the TLR signalling pathway (IκB) indicated activated gene expression when the snails were challenged with any of the used immune elicitors except wounding, although this was not seen in all individuals. Other components of the TLR pathway did not show a similar pattern. These findings suggest a general role of this pathway in controlling for the immune responses in our study species, which is in line with earlier findings in other organisms (Pila et al. 2016; Nie et al. 2018).

Manipulation of the environmental conditions also affected the expression levels of some of the annotated immune factors. Elevated temperature (25°C vs. 20°C) led to upregulation in cytokine MIF and antibacterial lysozymes. This result differs from earlier observations in which the effects of high temperature on antibacterial and PO-like activity of snail haemolymph have been negative at the phenotypic level (e.g., Seppälä and Jokela 2011; Leicht et al. 2017; Salo et al. 2017). In those studies, however, adverse effects were observed after a one-week exposure to high temperature (Leicht et al. 2013). In this study, the exposure lasted for two days. Together these findings suggest that the negative impacts of high temperature on snail immune function may only arise if the exposure time is long enough. Instead, short-term exposure to high temperature could have positive effects if it causes an increase in general performance, for example, owing to increased metabolic rate. In fact, an initial rise in snail growth rate and reproductive output under high temperature has been reported earlier (Leicht et al. 2013). Food deprivation had both positive and negative effects on the gene expression of certain components of the snail immune system (upregulation: MIF, LBP/BPI, lysozymes; downregulation: M-type lectins, glutathione). Earlier, food deprivation has been reported to reduce snail immune activity at the phenotypic level, although also this effect may depend on the duration of reduced food availability (Seppälä and Jokela 2010).

Gene expression of the heat shock proteins did not differ between control snails (20°C) and the individuals exposed to elevated temperature (25°C). This finding suggests that the high-temperature treatment may not have been highly stressful to snails. Although 25°C is above the thermal optimum of adult *L. stagnalis* snails (21-23°C depending on the trait) it was still not close to the critically high temperature of this species (Salo et al. 2019). Additionally, the exposure time was short compared to earlier studies (see the previous paragraph) and to a typical 8.4-day summer heatwave the snails are exposed to in nature (Meehl and Tebaldi 2004). Interestingly, gene expression of the heat shock proteins was reduced under food deprivation, which indicates their general role in stress responses and that the resource level of snails could affect their ability to tolerate exposure to high temperature. Such interactive effects should be examined in future studies. Furthermore, exposure of snails to both Gram-negative and Gram-positive bacteria increased the gene expression of ADH. This suggests that immune challenge can alter the metabolic rate of snails. However, although immune activation is often expected to increase metabolic activity (reviewed in Demas et al. 1997), other annotated factors with a potential role in snail metabolism were not affected by immune challenge treatments.

Our study was conducted using snails that originated from a genetically diverse laboratory stock that was formed by initially combining individuals from different natural populations (Langeloh et al. 2017). Genetically variable snails were used to identify genes that show high among-individual variation in gene expression. Such variation may arise, for example, from differences in genetic background and physiological condition of snails. Identifying genes that vary in their expression among individuals is very important for ecological and evolutionary studies that focus on understanding the sources and the consequences of trait variation among individuals. Understanding such variation can be critical because trait evolution may depend more strongly on variation in gene expression than differences in protein-coding sequences (Enard et al. 2002; Gilad et al. 2006). Moreover, the suitability of considering gene expression as a “trait”, and that it evolves in natural populations has been demonstrated in yeast (e.g., Ferea et al. 1999; Brem et al. 2002), fruit fly (e.g., Jin et al. 2001; Gibson et al. 2004), and fish (e.g., Oleksiak et al. 2005; Whitehead and Crawford 2006; Leder et al. 2015). Among-individual variation in gene expression is, however, easily overlooked in experiments that use genetically as homogeneous individuals as possible (e.g., one clone).

In our study, several components of the snail immune system from pathogen recognition (GNBP, lectins, TLR), to signalling/regulation (TLR pathway, cytokines, ROS production, apoptosis) and effectors (laccase, tyrosinase, antibacterial proteins) showed high variation in gene expression among snail individuals. In several cases, such variation was seen even among untreated snails (e.g., PGRP, TLR, Myd88, MIF, LBP/BPI, LAAOs, lysozymes, NOX, DUOX, TNF receptor). Two potential explanations are proposed for the latter finding. First, control snails may have harboured obscure infections from opportunistic microbes or viruses that activate snail immune function (Seppälä and Leicht 2013), leading to variation in transcription profiles. Second, these components of the snail immune system are expressed at a constant level within each individual, but the actual level varies among individuals (Bender et al. 2007). This would fit with innate type immunity, the expression levels reflecting the immediate capacity to respond to invading pathogens. Nevertheless, all the immune factors that showed high among-individual variation in expression levels are potentially interesting targets for future ecoimmunological research. To our knowledge, such variation has been utilised in ecoimmunology only in bumblebee (Brunner et al. 2013; Barribeau et al. 2014; Brunner et al. 2014; Marxer et al. 2016). In that species, condition dependence of immune defence and genetic specificity determining the outcome of a host-parasite interaction have been examined at the gene expression level. Similar variation may underlie the observation that parasite exposure does not yield infection in all snails of a generally susceptible strain in the *B. glabrata-S. mansoni* interaction (Théron and Coustau 2005).

Despite high among-individual variation in gene expression of many components of the snail immune system, our data did not reveal any individuals with generally very low or very high immune activity across the annotated factors. However, individuals within some of the immune activation treatments showed investment in different components of defence. For example, each snail exposed to *E. coli* showed high gene expression in two to four of the examined six antibacterial defence factors. Interestingly, which of these six factors showed high expression levels varied among individuals. Similarly, in snails exposed to trematode-infected snail tissue extracts, each individual typically showed high gene expression in only one, but not the same of the examined non-self recognition components (GNBP, FREP, galectins, TLR). Yet other immune properties vary among individuals both within field- and laboratory-maintained *L. stagnalis* populations. Van der Knaap et al. (1982; 1983) described two genetically-defined types of *L. stagnalis*, with “type I” snails having a particular type of lectin-mediated immune recognition that was absent in the majority of snails, designated as type II. These observations suggest that snails may show different immune defence strategies that lead to diverse responses even under a similar immune challenge. This calls for studies investigating among-individual variation in immune activity across immunological mechanisms in natural populations, as well as examining the contribution of such variation on snail performance (e.g., fitness components such as survival and fecundity).

In conclusion, our results confirm uniform aspects of gastropod/molluscan immunity, but also demonstrate apparent differences between *L. stagnalis* and some previously examined taxa. For instance, PRRs and lectins are present to detect pathogens and signal through the TLR pathway and cytokines to activate both cellular and humoral defence mechanisms. However, contrary to the high number of AMPs in other organisms, including molluscs, only a modest number of macin type AMPs was identified in *L. stagnalis*. Similarly, the low number of FREPs is comparable to *P. acuta* but differs from *B. glabrata* that shows high FREP diversity. Our data also indicate that various defence factors can contribute to the phenotypic immunological assays (antibacterial activity and PO-like activity of haemolymph) commonly used in earlier ecoimmunological research. Additionally, immune activation treatments (bacteria, trematodes) and manipulations of environmental conditions (ambient temperature, resource availability) revealed factors that contribute to snail immune responses (TLR pathway, FREP, cytokines, antibacterial defence, PO activity) and dependence of immune activity on environmental variation (cytokines, antibacterial defence, ROS production), as well as how these responses may depend on general stress reactions (HSP). Interestingly, many examined immune factors (PRRs, lectins, TLR pathway, cytokines, ROS production, apoptosis, PO enzymes, antibacterial proteins) showed considerable among-individual variation in the genetically diverse experimental snails used in this study. In addition to the components responding to immune elicitors, these factors can be important determinants of fitness variation in natural snail populations and should be included in future ecoimmunological studies.

ADH: alcohol dehydrogenase
AIF: apoptosis-inducing factor
AMP: antimicrobial peptide
CAT: catalase
DUOX: dual oxidase
DUOXA: oxidase maturation factor
ERR: estrogen-related receptor
FAIM1: Fas apoptotic inhibitory molecule 1
FBG: fibrinogen-like domain
FREP: fibrinogen-related protein
GNBP: Gram-negative bacteria binding-protein
GST: glutathione S-transferase
HSF: heat shock factor
HSP70: 70 kilodalton heat shock protein
HSP90: 90 kilodalton heat shock protein
HTRA2: serine protease
IAP: inhibitor of apoptosis proteins
IgSF: immunoglobulin superfamily
IKK: inhibitor of NF-κB kinase
IL-17: interleukin 17A
IκB: inhibitor of nuclear factor-κB
LAAO: L-amino acid oxidase
LBP/BPI: lipopolysaccharide-binding protein/bactericidal permeability-increasing protein
L-dopa: l-3,4-dihydroxyphenylalanine
LITAF: PS-induced TNF-α factor
LRR: leucine-rich repeat
MIF: macrophage migration inhibitory factor
MPEG: macrophage expressed gene
MyD88: myeloid differentiation primary response protein 88
NADPH: nicotinamide adenine dinucleotide phosphate
Nemo: NF-B essential modulator
NF-B: nuclear factor B
NF-κB: nuclear factor-κB
NOS: nitric oxide synthase
NOX: NADPH-oxidase
ORF: open reading frame
PARP: poly (ADP-ribose) polymerase
PC: principal component
PCA: principal component analysis
PGRP: peptidoglycan recognition protein
PO: phenoloxidase
PP1: protein phosphatase
PRR: pathogen-recognition receptors
RXR: retinoid acid receptor
SARM: sterile alpha and armadillo motif-containing protein
SOD and MnSOD: superoxidase dismutases
TAB: TAK1 binding protein
TAK1: transforming growth factor β-activated kinase-1
TIR: Toll-interleukin receptor
TLR: Toll-like receptor
TNF: tumor necrosis factor
TPM: transcripts per million
TRAF: tumor necrosis factor receptor-associated factor
TRF: TRAF-like protein
VIgLs: variable immunoglobulin and lectin-domain-containing molecules.

## Acknowledgements

We thank B. Hannah, S. Kobel, A. Minder, T. Torossi, and N. Zemp for help and guidance with the lab work and the analyses. We are grateful to M.-A. Coutellec, A. Dennis, B. Feldmeyer, J. Jokela, E. S. Loker, M. Neiman, and L. Shu for helpful discussions considering the experiment and the interpretation of the results. Data produced and analyzed in this paper were generated in collaboration with the Genetic Diversity Centre (GDC), ETH Zürich. The study was funded by the Emil Aaltonen Foundation and the Swiss National Science Foundation (grant 31003A 169531) to OS.

## Notes

### Competing Interest Statement

The authors have declared no competing interest.

