## Supplementary information for "Transcriptome profiling of a multiuse model species *Lymnaea stagnalis* (Gastropoda) for ecoimmunological research"

### Contents

**Table S1.** GenBank accession numbers of the sequences of proteins/genes relevant for immune function, stress responses, antioxidation and metabolism that were used as references in BLAST similarity searches to identify orthologs in the *L. stagnalis* reference transcriptome ........................ 3

**Table S2.** Summary of the annotated transcripts linked to proteins with a role in immune defence, stress-responses, or metabolism. The total number of identified transcripts, the number of transcripts with unique open reading frames (ORFs), an example domain architecture using SMART and PFAM domains[a] (when available in the annotation database), and transcripts representing unique ORFs for each examined factor are presented. Transcript ID the domain architecture refers to is in bold ........................................................................................ 7

**Table S1.** GenBank accession numbers of the sequences of proteins/genes relevant for immune function, stress responses, antioxidation and metabolism that were used as references in BLAST similarity searches to identify orthologs in the *L. stagnalis* reference transcriptome.

| Component | Reference(s) |
|---|---|
| **Non-self recognition** | |
| PGRP | AEH26026.1, AHB30456.1, XP_013064763.1, XP_013064406.1, XP_013062325.1, ABO40832.1, ABK76645.1, NP_001298227.1, XP_013063186.1, XP_013062465.1, XP_013062456.1, XP_005100262.1, XP_005093494.1, ABO40831.1, ABO40830.1, ABO40829.1 |
| GNBP | ABO40828.1 |
| FREP | Q8VCM7.1, ADA82222.1, CBM41040.1, AEO50745.1, O70165.1 |
| Galectin | ACO36044.1, NP_034837.2, ABQ09359.1, ACO53607.1, BAF75419.1 |
| Chi-lectin | O35744.2, P36222.2, Q1RQ16, Q1RQ22 |
| C-type lectins | AAX19697.1, ACS72237.1, AFA36633.1, ABO26594.1, P11226.2 |
| F-box lectins | Q96EF6.1, Q80UW2.1 |
| F-type lectins | Q7SIC1.1, ACF94293.1, ADJ40024.1, AFO64339.2, ADR63290.1 |
| I-type lectins | Q9Y286.1, AAH47244.1, AAA20148.1, P13597.1, Q91Y57.3 |
| L-type lectin | P49257.2, Q12907.1, EKC31192.1, EKC20002.1 |
| M-type lectin | Q925U4.1, Q91VV3, EKC26076.1, EKC23948.1 |
| P-type lectins | CAA45423.1, EKC18244.1, NP_000867.2, EKC35005.1 |
| R-type lectins | BAA36393.1, BAL61208.1, Q2HZ94 |
| Intelectins | ACC62155.1, CAJ32677.2, ACC62157.1, ACC62156.1, NP_001082570.1 |
| Pentraxins | AAZ68032.1, ABG37054.1, P02743.2, P02741.1 |
| **TLR pathway** | |
| TLR | AGB93809.1 |
| MyD88 | XP_013086371.1, XP_013086405.1 |
| IRAK | XP_013088618.1 |
| TRAF | XP_013069680.1, XP_013069680.1 |
| TBK | XP_013083625.1 |
| TAK | XP_013073733.1, XP_013073731.1, XP_013073732.1, XP_013073736.1, XP_013073734.1, XP_013073735.1 |
| TRF | XP_013089600.1, XP_013073636.1, XP_005088853.1, XP_005104410.1, XP_012934632.1 |
| TAB | XP_013096269.1, XP_013096268.1, XP_005094474.1, XP_005094473.1, XP_014776667.1 |
| SARM | XP_013080872.1, XP_013080871.1 |

| | |
|---|---|
| IKKα | XP_005096610.2 |
| IKKγ Nemo | XP_013073873.1 |
| LITAF | XP_013089264.1, XP_013089261.1, XP_013089265.1, XP_013089270.1, XP_013089262.1, XP_013089269.1 |
| IκB | NP_001298198.1 |
| NF-κBp65 | NP_001298191.1, ACN73461.1 |
| NF-κBp100 | XP_022318641.1, XP_022318640.1 |
| MPEG | XP_013065342.1 |
| **Cytokines** | |
| MIF | NP_001298224.1, XP_013075404.1, XP_005103629.2, XP_012939653.1, AQY19126.1 |
| IL17 | XP_013096467.1, XP_013090467.1, XP_013090466.1, XP_013085850.1, XP_013085675.1, XP_013080675.1, XP_013065647.1, XP_021365521.1, AGZ03660.1, XP_021365518.1, XP_009058928.1, XP_021365519.1, XP_022315060.1, XP_022310682.1, AKM49935.1, XP_009048356.1, XP_005097238.1, XP_011438620.1, NP_001292220.1, EKC33705.1, AKM49938.1, XP_022286467.1, XP_021365533.1, XP_022344404.1, XP_022344403.1, NP_001295831.1, OWF44912.1, OWF56800.1, XP_021364761.1, XP_011438445.1, AKQ98198.1, XP_009051888.1, NP_001295781.1, AKQ98197.1, XP_011414373.1, XP_022318598.1, XP_022315342.1 |
| TNF | XP_013076951.1, XP_013061250.1, XP_013061249.1, XP_013061248.1, XP_013061247.1, XP_013096914.1, XP_013096721.1, XP_013096720.1, XP_013096719.1, XP_013096712.1, XP_013096711.1, XP_013096710.1, XP_013094785.1, XP_013094784.1, XP_013094783.1, XP_013094495.1, XP_013094494.1, XP_013086591.1, XP_013082590.1, XP_013082589.1, XP_013082588.1, XP_013082587.1, XP_013076957.1, XP_013073494., XP_013073185.1, XP_013072204.1, XP_013072202.1, XP_013070598.1, XP_013067540.1, XP_013067539.1, XP_013067535.1, XP_013070598.1, XP_013077259.1, XP_022338094.1, XP_022338095.1, XP_011450914.1, EKC35160.1, XP_021346927.1, XP_021364181.1, XP_013073494.1, XP_012936811.1, XP_021346926.1, OWF56727.1, XP_021340535.1 |
| **Antimicrobial defence** | |
| Macins | AFR36920.1, CK989857.1 |
| LBP/BPI | AGG82435.1, AKM45823.1, AKM45822.1, AKM45821.1, AKM45820.1 |
| LAAO | NP_001191524.1, AAN78211.1, Q17043.1, P35903.1, AAR14185.1, AAR14187.1, AAR14186.1 |
| Lysozymes | AGQ50333.1, AGQ50332.1, AGQ50331.1, ADR70996.1, ADR70995.1, AGQ50337.1, AGQ50335.1, AGQ50330.1, ADV36303.1, AGQ50336.1, AGQ50334.1, AOX15710.1, AOX15709.1, AOX15708.1, AOX15707.1, XM_013226064.1 |

| | |
|---|---|
| Biomphalysin | AGG38744.1, NP_001298219.1 |
| Glabralysin | A0A2C9L301, A0A2C9M7U6, A0A2C9M7H5, A0A2C9L4Y1, A0A2C9LNR7, A0A2C9KPA0 |
| **Production of reactive oxygen species** | |
| NOX | XP_005090645.1, XP_005096802.1, XP_012938230.1 |
| DUOX | XP_012942535.1, XP_012943483.1 |
| DUOXA | XP_021362528.1, XP_022343917.1 |
| NOS | NP_001191470.1, AGI44587.1 |
| Peroxidasin | XP_013088968.1, XP_005110224.1 |
| Glutaredoxin | XP_013084576.1, XP_013061972.1, XP_005104531.1, XP_005110052.1 |
| GST | XP_013075526.1, XP_012944480.1, XP_013071551.1, XP_013060671.1, XP_012945427.1, RUS76363.1, XP_013097111.1, AEI27296.1, XP_005109181.1, XP_013070482.1, AXN72681.1, XP_013095077.1, XP_009049471.1, RUS84494.1, XP_005094586.1, XP_005106935.1, XP_012941915.1 |
| Nucleoredoxin | EKC27452.1, XP_013073587.1, XP_005098394.1, XP_013074893.1 |
| Peroxiredoxin | XP_013086205.1, XP_013087397.1, XP_013091924.1, ACI42883.1 |
| **Phenoloxidase/melanisation-type reaction** | |
| proPO | CCQ18551.1, NP_476812.1, AGC54940.1, BAF98646.1 |
| Laccase | XP_013079632.1, XP_013079631.1, XP_013086277.1, XP_013074984.1, XP_013074983.1, XP_013074982.1, XP_013068526.1, XP_013085808.1, XP_013062555.1, XP_013064545.1, XP_013079036.1, XP_013085832.1, XP_013085833.1, XP_013085834.1, XP_013088628.1, XP_013090126.1, XP_013067361.1, XP_013067388.1, XP_013074978.1, XP_013074979.1, XP_013074980.1, XP_013074981.1, ESO98873.1, ESO97324.1, ESO96136.1, ESO89911.1, XP_009059385.1, XP_009053251.1, XP_009051926.1, XP_009050501.1, ESO89918.1, XP_009059392.1, ESO90420.1, ESO89919.1, ESO84603.1, XP_009064708.1, XP_009059393.1, XP_009058892.1, XP_012937904.1, XP_012944262.1, XP_012942999.1, XP_012942636.1, XP_012940646.1, XP_012945653.1, XP_012934797.1 |
| Tyrosinase | XP_013080599.1, XP_013065857.1, APC92582.1, EKC35331.1, EKC38463.1, OWF48254.1, APC92581.1, XP_021374137.1, AMB26746.1, ASR73340.1, ALG64484.1 |
| **Apoptosis** | |
| FAIM1 | XP_013071009.1 |
| HTRA2 | XP_013072351.1 |
| AIF | XP_013095497.1, XP_013095498.1 |
| IAP | XP_013075317.1, XP_013064429.1, XP_013064681.1, XP_013075313.1, XP_013060630.1, XP_013090049.1, XP_013064690.1, XP_013075320.1, XP_013082134.1, XP_013084114.1, XP_013084111.1 |

| | |
|---|---|
| Bcl-Bax | XP_013075909.1, XP_013089117.1 |
| PARP | XP_013090488.1, XP_013062683.1, XP_013062670.1, XP_013065538.1, XP_013068782.1, XP_013062696.1, XP_013067682.1, XP_013092390.1, XP_013096616.1 |
| Caspase | XP_013067788.1, XP_013065656.1, XP_013073999.1, XP_013074008.1, XP_013088924.1, XP_013094549.1, XP_013095352.1 |
| **Stress responses** | |
| HSP70 | XP_013081556.1, XP_013082114.1, ABB45831.1, XP_013091813.1 |
| HSP90 | XP_013072750.1 |
| HSF | XP_013085217.1 |
| **Antioxidant enzymes** | |
| SOD | AAP93637.2, XP_013070344.1 |
| MnSOD | NP_001298192.1 |
| CAT | ACO59957.1 |
| Glutathione peroxidase | ACO59958.1 |
| Glutathione reductase | ACO59956.1 |
| **Metabolism** | |
| PP1 | XP_013085822.1 |
| Ubiquitin | NP_001298238.1 |
| Ferritin | XP_013080834.1 |
| ADH | XP_013073559.1 |
| ERR | XP_013094385.1, XP_013080351.1 |
| RXR | NP_001298239.1 |

**Table S2.** Summary of the annotated transcripts linked to proteins with a role in immune defence, stress-responses, or metabolism. The total number of identified transcripts, the number of transcripts with unique open reading frames (ORFs), an example domain architecture using SMART and PFAM domains[a] (when available in the annotation database), and transcripts representing unique ORFs for each examined factor are presented. Transcript ID the domain architecture refers to is in bold.

| Component | # transcripts | # unique ORFs | Example of domain structure | Transcript IDs with unique ORFs |
|---|---|---|---|---|
| **Non-self recognition** | | | | |
| PGRPs | 8 | 7 | 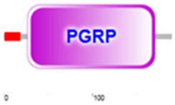 | 14145_c0_g1_i1<br>**32865_c0_g1_i1**<br>34108_c0_g1_i1<br>36537_c0_g1_i1<br>36963_c0_g1_i1<br>42094_c0_g6_i1<br>42956_c0_g1_i1 |
| GNBPs | 8 | 4 | 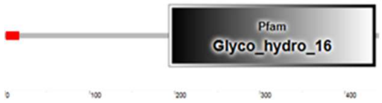 | **41774_c0_g1_i2**<br>47227_c0_g2_i1<br>47391_c1_g1_i1<br>48085_c1_g8_i11 |
| FREPs | 2 | 2 | 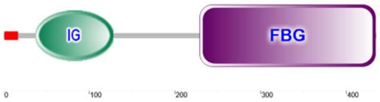 | **36764_c0_g1_i2**<br>45250_c0_g1_i1 |
| Galectins | 20 | 6 | 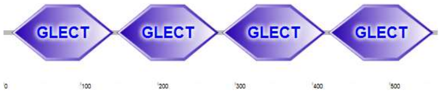 | 42521_c0_g1_i2<br>44043_c0_g3_i1<br>**44175_c0_g1_i1**<br>47859_c0_g2_i1<br>47859_c0_g2_i5<br>48059_c1_g2_i1 |
| Chi-lectins | 11 | 8 | 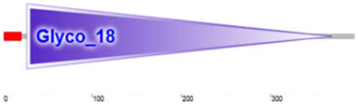 | **33218_c0_g1_i1**<br>44081_c0_g1_i1<br>44227_c0_g2_i2<br>45634_c0_g3_i1<br>47306_c0_g5_i1<br>48292_c1_g1_i3<br>26259_c0_g1_i1<br>45255_c1_g3_i3 |
| L-type lectins | 3 | 2 | 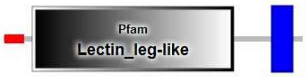 | **44959_c0_g1_i1**<br>48268_c2_g1_i1 |
| M-type lectins | 9 | 3 | 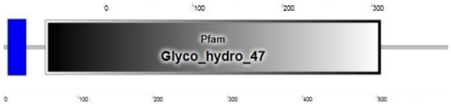 | 37655_c0_g1_i1<br>39342_c0_g1_i1<br>**47242_c4_g2_i1** |
| **TLR pathway** | | | | |
| TLR | 1 | 1 | 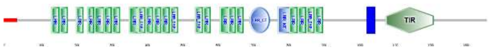 | 28651_c0_g1_i1 |

| | | | | |
|---|---|---|---|---|
| MyD88 | 7 | 1 | 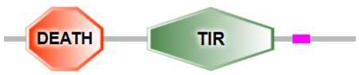 | 47065_c0_g1_i1 |
| TRAF | 1 | 1 | 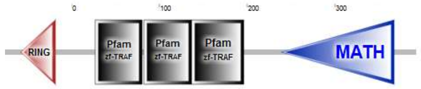 | 22931_c0_g1_i1 |
| TRF | 1 | 1 | 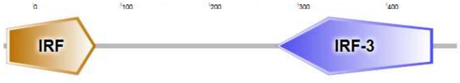 | 34891_c0_g1_i1 |
| TAB | 1 | 1 | 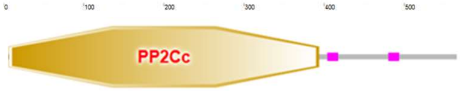 | 24669_c0_g1_i1 |
| SARM | 1 | 1 | 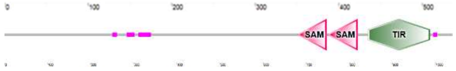 | 46010_c7_g2_i1 |
| IKKα | 2 | 1 | 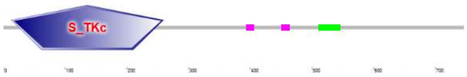 | 35196_c0_g1_i1 |
| IKKγ Nemo | 6 | 1 | 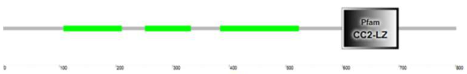 | 47446_c1_g1_i1 |
| LITAF | 6 | 2 | 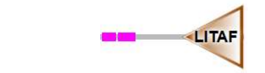 | **45095_c0_g1_i1**<br>46794_c1_g1_i1 |
| IκB | 2 | 2 | 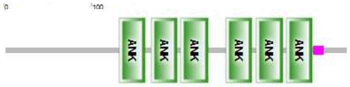 | **39554_c0_g1_i1**<br>46270_c2_g2_i1 |
| NF-κBp65 | 4 | 2 | 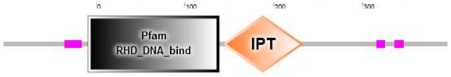 | 38589_c1_g1_i1<br>**43263_c0_g2_i1** |
| MPEG | 3 | 2 | 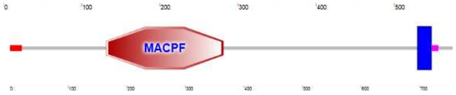 | **46200_c0_g1_i1**<br>47480_c0_g1_i1 |
| **Cytokines** | | | | |
| MIF | 2 | 1 | 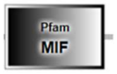 | 36453_c0_g1_i1 |
| IL-17 | 3 | 3 | 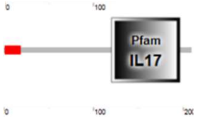 | **110902_c0_g1_i1**<br>4951_c0_g1_i1<br>17215_c1_g1_i1 |
| TNF | 63 | 23 | 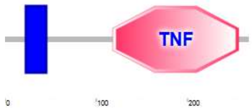 | 15848_c0_g1_i1<br>**20877_c0_g2_i1**<br>34263_c0_g1_i2<br>37129_c0_g1_i1<br>37407_c0_g1_i1<br>37593_c0_g1_i1<br>39166_c0_g1_i1<br>41927_c0_g1_i1<br>42168_c1_g1_i2<br>42302_c0_g4_i1<br>42612_c1_g1_i1<br>42736_c0_g1_i1<br>43123_c0_g2_i2 |

| | | | | |
|---|---|---|---|---|
| | | | | 43870_c0_g1_i2<br>44512_c0_g1_i2<br>45558_c0_g3_i1<br>45927_c0_g1_i1<br>46059_c2_g1_i2<br>46501_c1_g2_i1<br>47286_c3_g3_i2<br>47468_c1_g1_i1<br>47690_c7_g2_i10<br>48316_c0_g1_i1 |
| **Antimicrobial defence** | | | | |
| Macins | 7 | 6 | 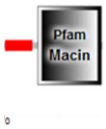  | **18198_c0_g1_i1**<br>18890_c0_g1_i1<br>34989_c0_g2_i1<br>35624_c0_g1_i1<br>41887_c0_g3_i1<br>43018_c0_g6_i3 |
| LBP/BPI | 4 | 3 | 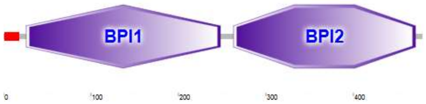  | **25950_c0_g1_i1**<br>45368_c0_g6_i1<br>46649_c0_g1_i1 |
| LAAO | 6 | 3 | 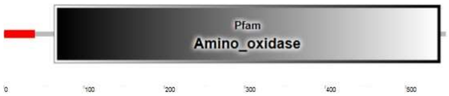  | **47821_c4_g3_i1**<br>47893_c0_g9_i1<br>48351_c1_g3_i1 |
| Lysozymes | 11 | 4 | | 28373_c0_g1_i1<br>35313_c1_g1_i1<br>36143_c0_g3_i1<br>44233_c1_g1_i1 |
| Lymnaea-lysins | 4 | 3 | 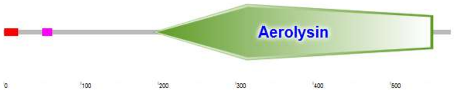  | **38389_c0_g1_i1**<br>47349_c1_g38_i2<br>5994_c0_g2_i1 |
| Stagnalysins | 3 | 3 | | 34104_c0_g1_i1<br>42332_c1_g1_i1<br>48069_c5_g1_i1 |
| **Production of reactive oxygen species** | | | | |
| NOX | 9 | 3 | 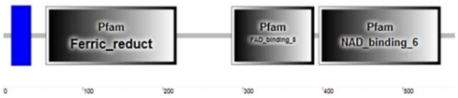  | **40220_c1_g2_i1**<br>36496_c0_g1_i1<br>36278_c0_g1_i1 |
| DUOX | 6 | 2 | 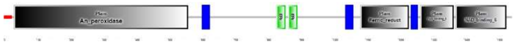 | **42865_c0_g1_i3**<br>46890_c0_g1_i1 |
| DUOXA | 1 | 1 | 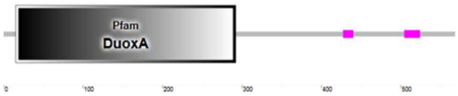  | 38205_c0_g1_i1 |
| NOS | 3 | 1 | 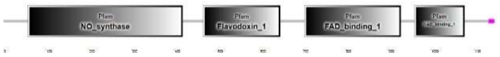 | 45570_c1_g1_i1 |
| Peroxidasin | 16 | 6 | | 40499_c0_g1_i1<br>45939_c0_g1_i1 |

| | | | | |
|---|---|---|---|---|
| | | | | 46236_c0_g1_i2<br>46303_c1_g1_i1<br>47256_c0_g1_i1<br>48181_c2_g1_i1 |
| Glutaredoxin | 7 | 4 | 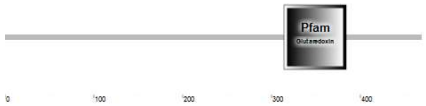 | **34293_c0_g2_i1**<br>34343_c1_g2_i1<br>4572_c0_g2_i1<br>45783_c0_g1_i1 |
| GST | 32 | 17 | 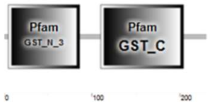 | **1429_c0_g2_i1**<br>31273_c0_g1_i1<br>36750_c0_g1<br>38055_c0_g1_i2<br>39155_c0_g1_i1<br>40581_c0_g1_i1<br>40585_c0_g1_i1<br>41000_c0_g2_i1<br>43131_c0_g1_i1<br>44207_c5_g1_i1<br>44723_c0_g5_i1<br>46105_c0_g60_i1<br>46670_c0_g1_i1<br>46742_c1_g1_i1<br>46851_c0_g1_i2<br>46851_c0_g1_i6<br>46929_c1_g35_i1 |
| Nucleoredoxin | 7 | 4 | 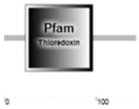 | **33046_c0_g1_i1**<br>39371_c0_g1_i1<br>44859_c0_g2_i1<br>45653_c0_g1_i2 |
| Peroxiredoxin | 9 | 5 | 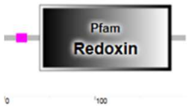 | **27522_c0_g1_i1**<br>34959_c0_g2_i1<br>41634_c2_g1_i1<br>46977_c0_g1_i1<br>47425_c0_g2_i6 |
| **Phenoloxidase/melanisation** | | | | |
| Laccase | 19 | 8 | 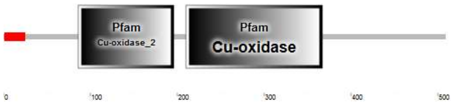 | **35966_c0_g1_i1**<br>38462_c0_g1_i1<br>45077_c0_g1_i1<br>47321_c2_g2_i1<br>47844_c2_g5_i1<br>48217_c1_g1_i10<br>38987_c2_g2_i1<br>45762_c4_g1_i3 |
| Tyrosinase | 2 | 2 | 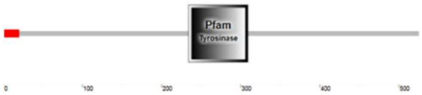 | **26246_c2_g1_i1**<br>46270_c3_g2_i1 |

| | | | | |
|---|---|---|---|---|
| **Apoptosis** | | | | |
| FAIM1 | 1 | 1 | 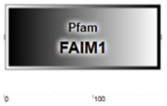 | 44771_c0_g1_i1 |
| HTRA2 | 3 | 2 | 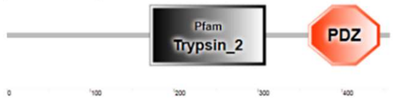 | **36021_c0_g1_i1**<br>37936_c0_g1_i1 |
| AIF | 1 | 1 | 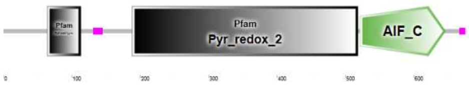 | 40505_c0_g1_i1 |
| IAP | 25 | 10 | 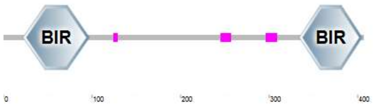 | **30243_c0_g1_i1**<br>41110_c0_g1_i1<br>42472_c1_g1_i1<br>46127_c2_g1_i1<br>47200_c0_g1_i1<br>47310_c1_g1_i4<br>47642_c0_g1_i11<br>48019_c3_g3_i1<br>48046_c0_g1_i3<br>48048_c3_g1_i2 |
| Bcl-Bax | 1 | 1 | 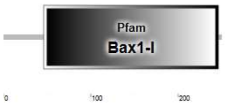 | 44934_c1_g1_i3 |
| PARP | 21 | [b] | | |
| Caspase | 20 | 8 | 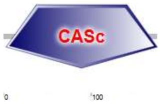 | **33457_c0_g1_i1**<br>37005_c0_g1_i1<br>37479_c0_g2_i1<br>42025_c0_g1_i2<br>43539_c0_g1_i1<br>45052_c4_g2_i1<br>48203_c0_g1_i2<br>48216_c2_g3_i10 |
| **Stress responses** | | | | |
| HSP70 | 13 | 5 | 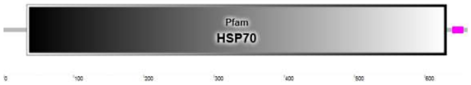 | **40510_c1_g2_i1**<br>41463_c0_g1_i1<br>44096_c1_g1_i1<br>46830_c1_g3_i1<br>47977_c0_g5_i1 |
| HSP90 | 1 | 1 | 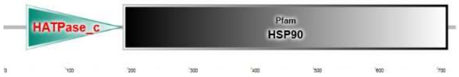 | 47161_c0_g4_i1 |
| HSF | 1 | 1 | 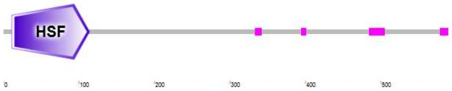 | 29117_c0_g1_i1 |
| **Antioxidant enzymes** | | | | |
| SOD | 6 | 2 | 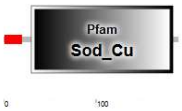 | **41920_c0_g1_i2**<br>45899_c0_g2_i1 |

| | | | | |
|---|---|---|---|---|
| MnSOD | 1 | 1 | 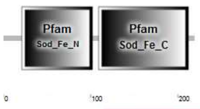 | 28257_c0_g1_i1 |
| CAT | 2 | 2 | 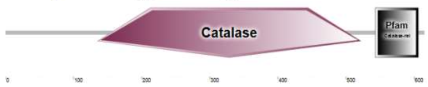 | **40830_c0_g1_i1**<br>44469_c0_g1_i1 |
| glutathione peroxidase | 5 | 3 | 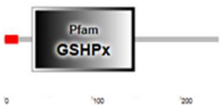 | **36171_c0_g2_i1**<br>37235_c0_g1_i1<br>47906_c1_g7_i1 |
| glutathione reductase | 1 | 1 | 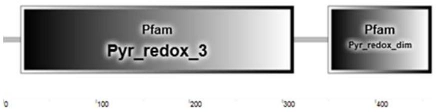 | 44983_c1_g1_i1 |
| **Metabolism** | | | | |
| PP1 | 2 | 1 | 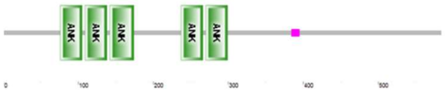 | 36580_c0_g1_i1 |
| Ubiquitin | 13 | 8 | 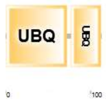 | **109816_c1_g1_i1**<br>33535_c0_g1_i1<br>40537_c0_g1_i1<br>44027_c1_g1_i2<br>44915_c1_g1_i1<br>45261_c0_g1_i1<br>46234_c4_g1_i1<br>46544_c1_g3_i2 |
| Ferritin | 3 | 3 | 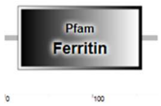 | **125081_c0_g1_i1**<br>33209_c0_g1_i1<br>43780_c0_g5_i1 |
| ADH | 3 | 1 | 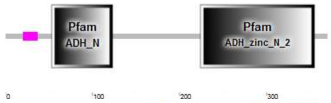 | 47996_c0_g14_i4 |
| ERR | 2 | 2 | 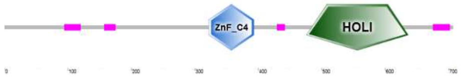 | **36130_c1_g1_i4**<br>42943_c0_g1_i1 |
| RXR | 5 | 3 | 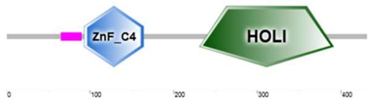 | 33793_c0_g1_i1<br>41983_c1_g1_i1<br>42786_c1_g1_i1 |

[a]SMART/PFAM domains: ADH_N – alcohol dehydrogenase GroES-like domain; ADH_zinc_N_2 – zinc-binding dehydrogenase; Aerolysin – aerolysin toxin; AIF_C – apoptosis-inducing factor, mitochondrion-associated, C-term; Amino_oxidase – flavin-containing amine oxidoreductase; An_peroxidase – animal heme-dependent peroxidase; An_peroxidase – animal heme-dependent peroxidase; ANK – ankyrin repeats; Bax1-I – inhibitor of apoptosis-promoting Bax1; BIR – baculoviral inhibition of apoptosis protein repeat; BPI1 – bactericidal permeability-increasing protein / lipopolysaccharide-binding protein / cholesteryl ester transfer protein N-terminal domain; BPI2 – bactericidal permeability-increasing protein / lipopolysaccharide-binding protein / cholesteryl ester transfer protein C-terminal domain; CASc – caspase, interleukin-1 beta converting enzyme homologues; Catalase – catalase for enzymatic detoxification of hydrogen peroxide; CC2-LZ – leucine zipper of domain CC2 of NEMO, NF-kappa-B essential modulator; Cu-oxidase – multicopper oxidase; DEATH – DEATH domain, bind each other to form oligomers of factors

involved in signaling pathways, frequently associated with immunity or apopotosis; Deltameth_res – deltamethrin resistance; DuoxA – dual oxidase maturation factor; EHf – EF-hand, calcium binding motif; FAD_binding_1 – flavoprotein pyridine nucleotide cytochrome reductase; FAD_binding_8 – FAD binding domain associated with ferric reductase NAD binding proteins and the heavy chain of Cytochrome b-245; FAD_binding_8 – FAD-binding domain; FAIM1 – Fas apoptotic inhibitory molecule; FBG – fibrinogen-like domain; Ferric_reduct – ferric reductase like transmembrane component; Ferritin – ferritin-like domain; Flavodoxin_1 – flavodoxin-like domain; GLECT – galectin; Glutaredoxin – glutaredoxin, oxidation repair enzyme; Glyco_18 – O-Glycosyl hydrolases; Glyco_hydro_16 – glycosyl hydrolases family 16; Glyco_hydro_47 – glycoside hydrolase family 47; GSHPx – glutathione peroxidase; GST_C – glutathione S-transferase, C-terminal domain; GST_N_3 – glutathione S-transferase; HATPase_c – histidine kinase-like ATPases; HOLI – ligand binding domain of hormone receptors; HSF – heat shock factor; HSP70 – heat shock protein 70; HSP90 – heat shock protein 90; IG – immunoglobulin; IL17 – interleukin-17; IPT – ig-like, plexins, transcription factors; IRF – interferon-regulatory factor; IRF-3 – interferon-regulatory factor 3; Lectin_leg-like – L-type lectin domain; LITAF – LPS-induced TNF-activating factor; LRR – leucine-rich repeats; Macin – macin family; MACPF – membrane-attack complex / perforin; MATH – meprin and TRAF homology; MIF – macrophage migration inhibitory factor; NAD_binding_1 – oxidoreductase NAD-binding domain; NAD_binding_6 – ferric reductase NAD binding domain; NO_synthase – nitric oxide synthase; PBRP – animal peptidoglycan recognition protein; PDZ – domain present in PSD-95, Dlg, and ZO-1/2, proteins that anchor receptor proteins to the cytoskeleton, to help form signal transduction complexes; PP2Cc – serine/threonine phosphatases family 2C, catalytic domain; Pyr_redox_2 – pyridine nucleotide-disulphide oxidoreductase; Pyr_redox_3 – pyridine nucleotide-disulphide oxidoreductase; Pyr_redox_dim – pyridine nucleotide-disulphide oxidoreductase, dimerisation domain; Redoxin – redoxin domain with oxidoreductase activity; RHD_DNA_bind – Rel homology DNA-binding domain; RING – ring finger; S_TKc – serine/Threonine protein kinases, catalytic domain; SAM – sterile alpha motif; Sod_Cu & Sod_Fe_N & Sod_Fe_C – catergories of superoxide dismutase; Thioredoxin – thioredoxin, small redox proteins involved in responses to reactive oxygen species and cell-cell communication; TIR – Toll/interleukin-1 receptor; TNF – tumour necrosis factor family; Trypsin_2 – trypsin-like peptidase domain; Tyrosinase – tyrosinase, enzymatic domain of copper-type 3 enzymes involved in melanization contributing to pigmentation or immune defence; UBQ – ubiquitin homologues; zf-TRAF – TRAF-type zinc finger; ZnF_C4 – c4 zinc finger in nuclear hormone receptors.

[b]Determining which sequences cover a complete ORF not possible.

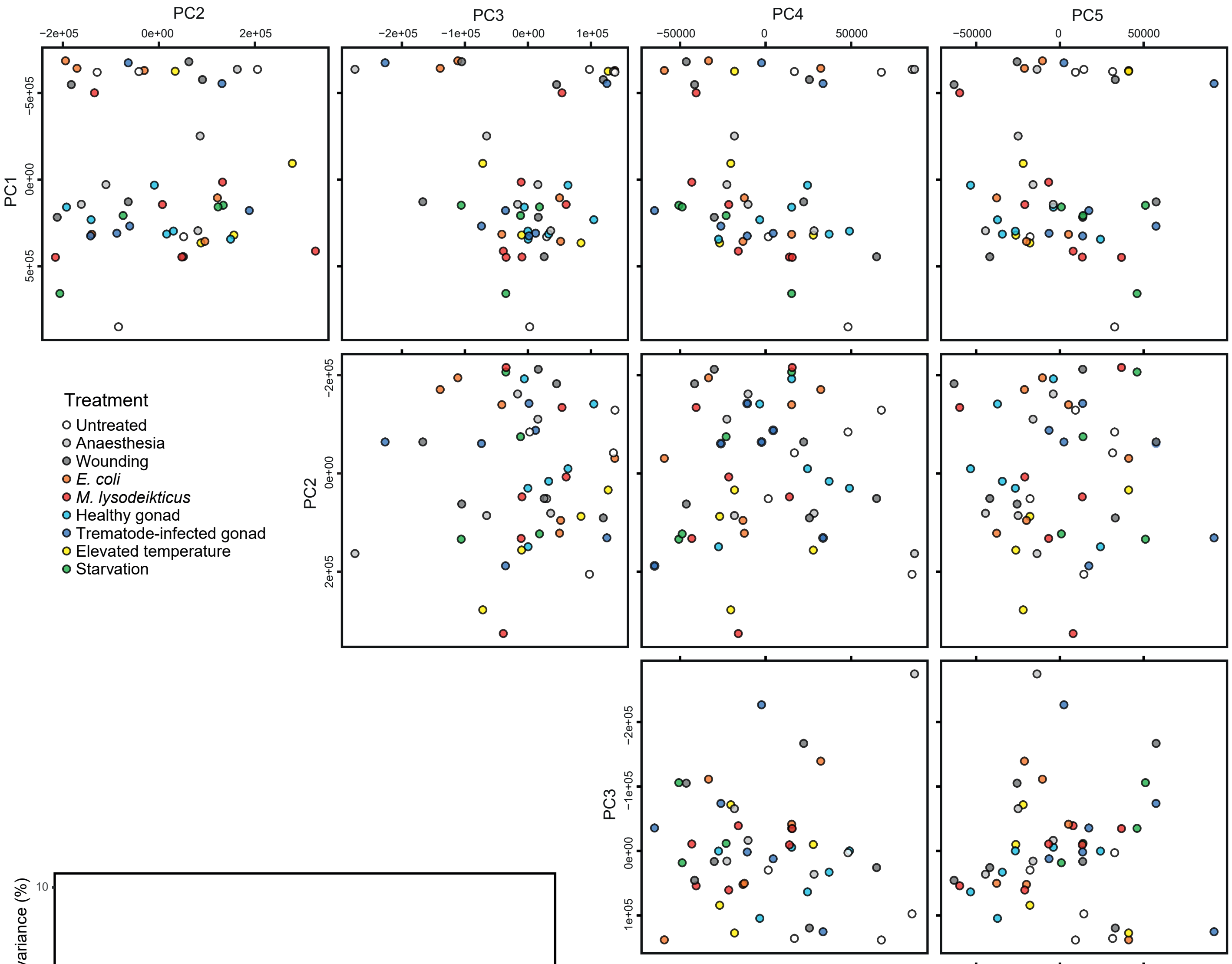

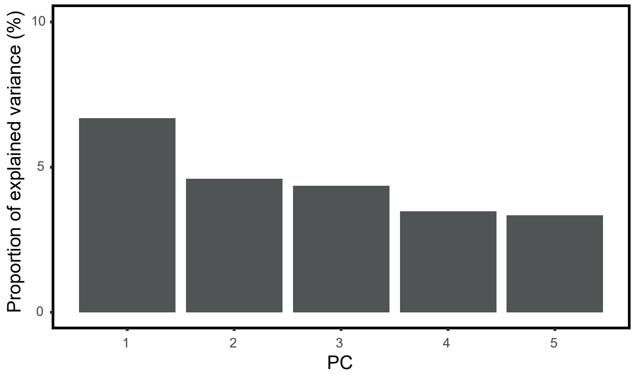

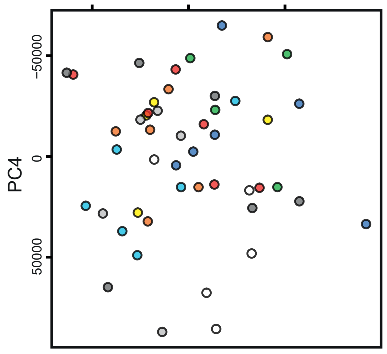

Fig. S1. Principal component analysis (PCA) plots showing the variation in transcriptome-wide expression profiles of the experimental snails using the first five principal components (PCs) after internal normalization in Sleuth. Additionally, the proportion of total variance each PC explained in the data is presented.

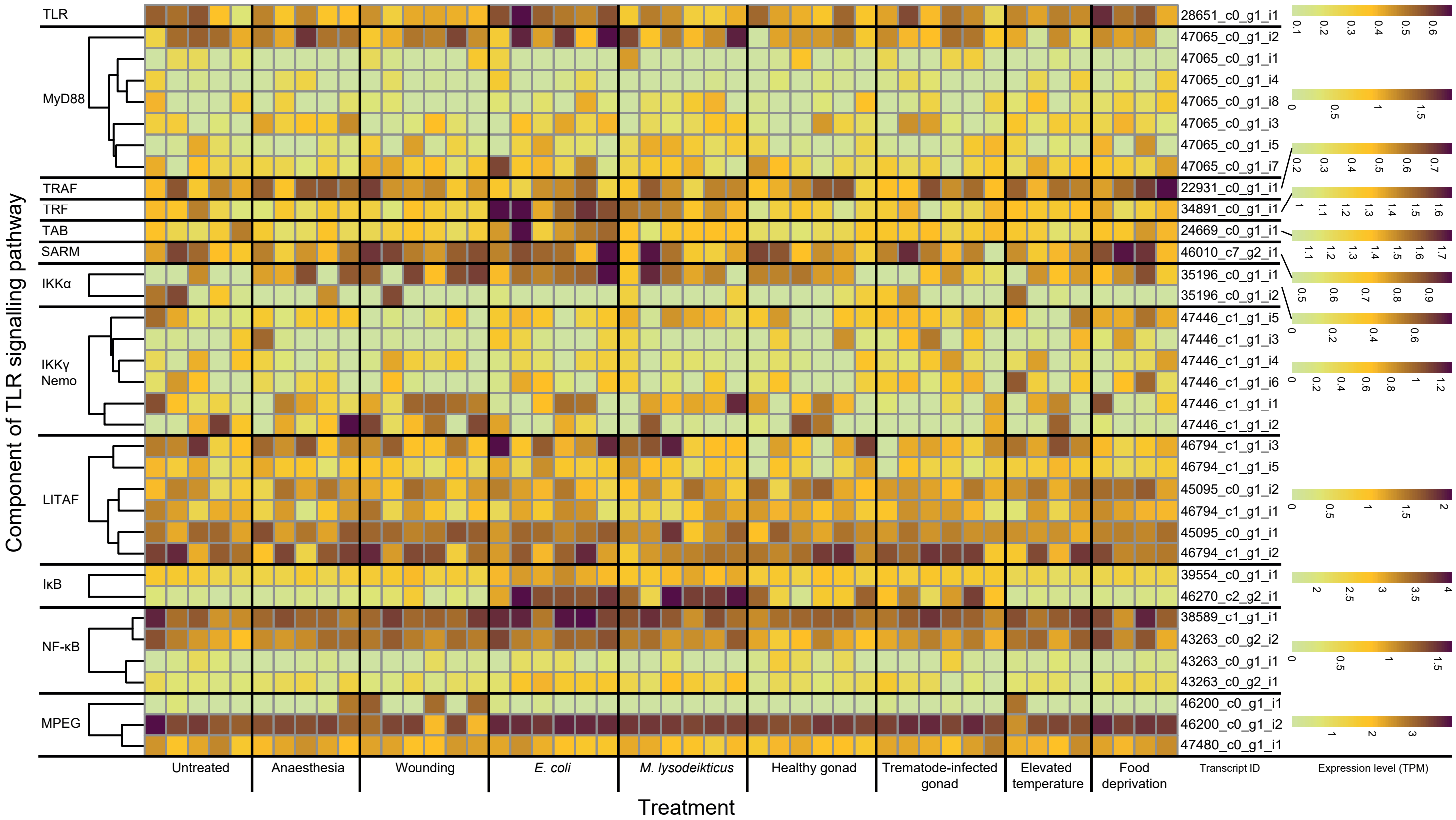

Fig. S2. Expression levels of individual transcripts found to represent annotated factors related to TLR signalling pathway in units of transcripts per million (TPM) for each experimental snail. Heatmap shows the variation for each factor using the dynamic range. Transcripts related to each factor are clustered according to their similarity.

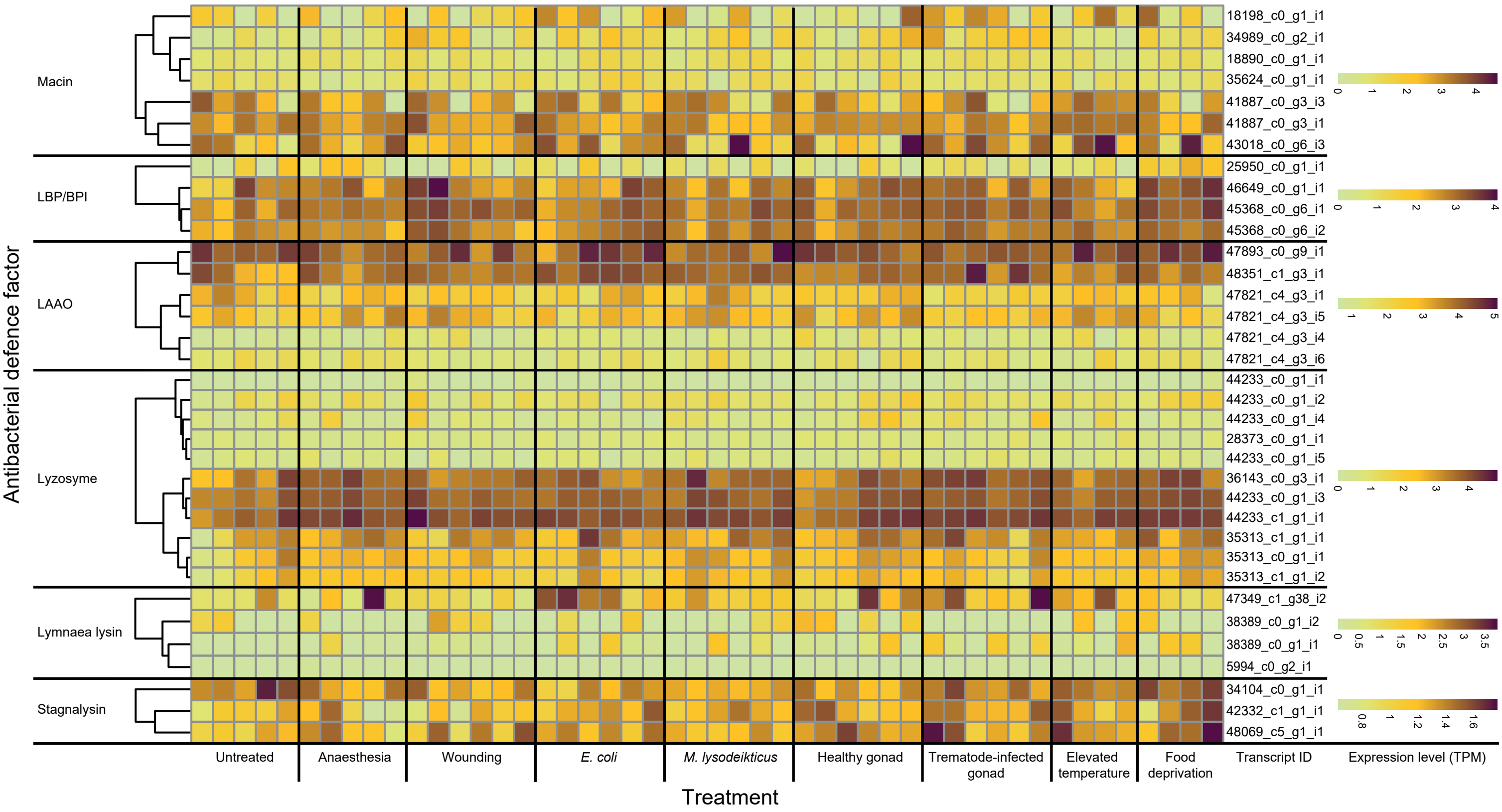

Fig. S3. Expression levels of individual transcripts found to represent the annotated antibacterial defence factors in units of transcripts per million (TPM) for each experimental snail. Heatmap shows the variation for each factor using the dynamic range. Transcripts related to each factor are clustered according to their similarity.

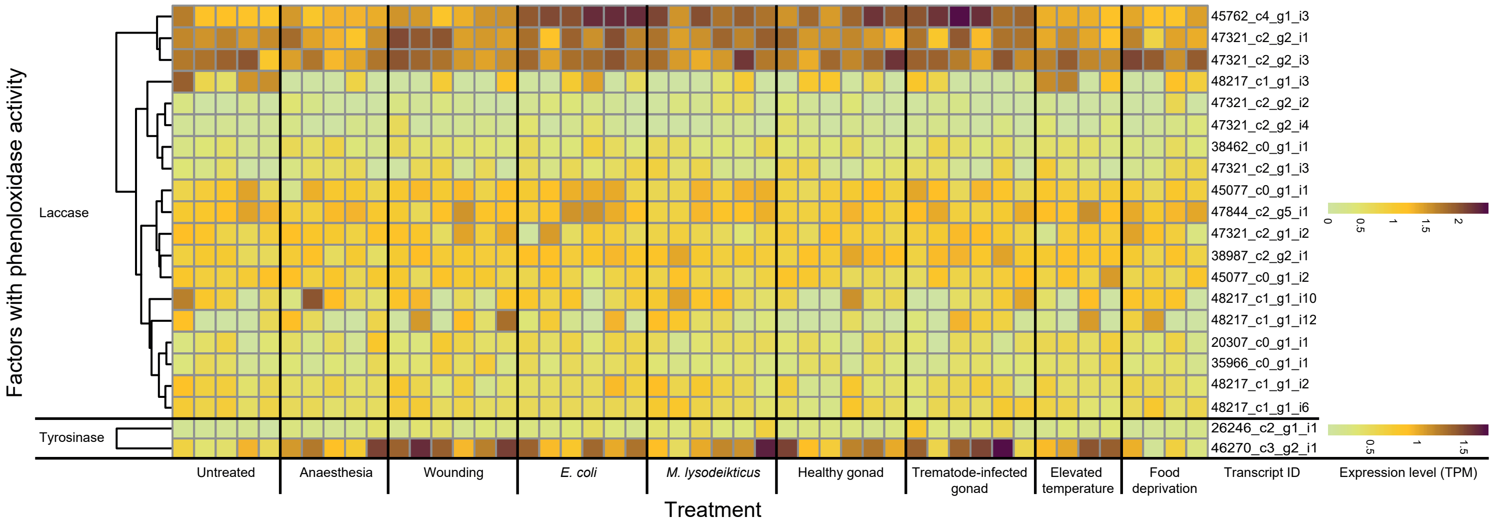

Fig. S4. Expression levels of individual transcripts found to represent annotated factors related to phenoloxidase/melanisation-type reaction in units of transcripts per million (TPM) for each experimental snail. Heatmap shows the variation for each factor using its dynamic range. Transcripts related to each factor are clustered according to their similarity.

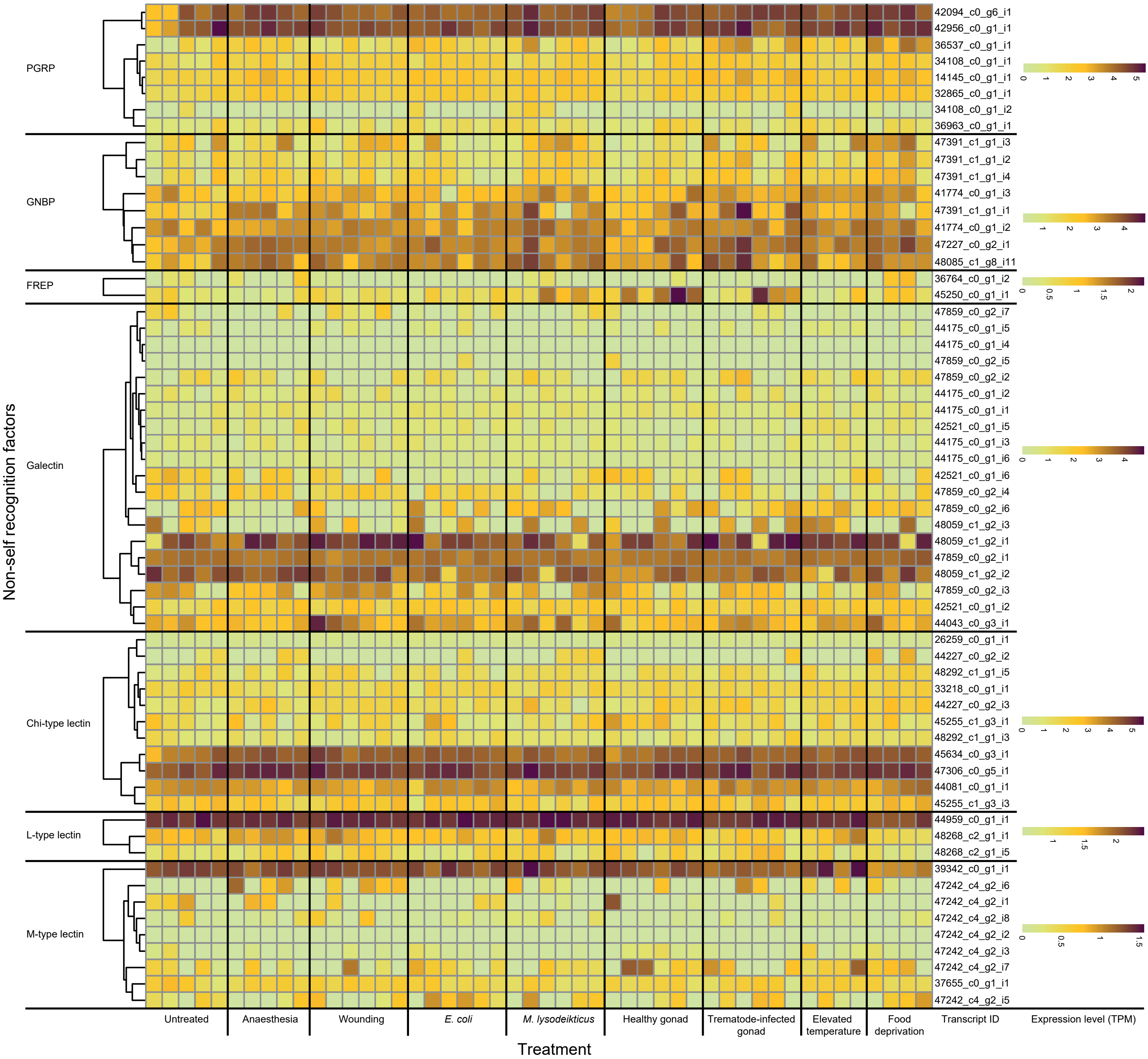

Fig. S5. Expression levels of individual transcripts found to represent annotated factors related to non-self recognition in units of transcripts per million (TPM) for each experimental snail. Heatmap shows the variation for each factor using the dynamic range. Transcripts related to each factor are clustered according to their similarity.

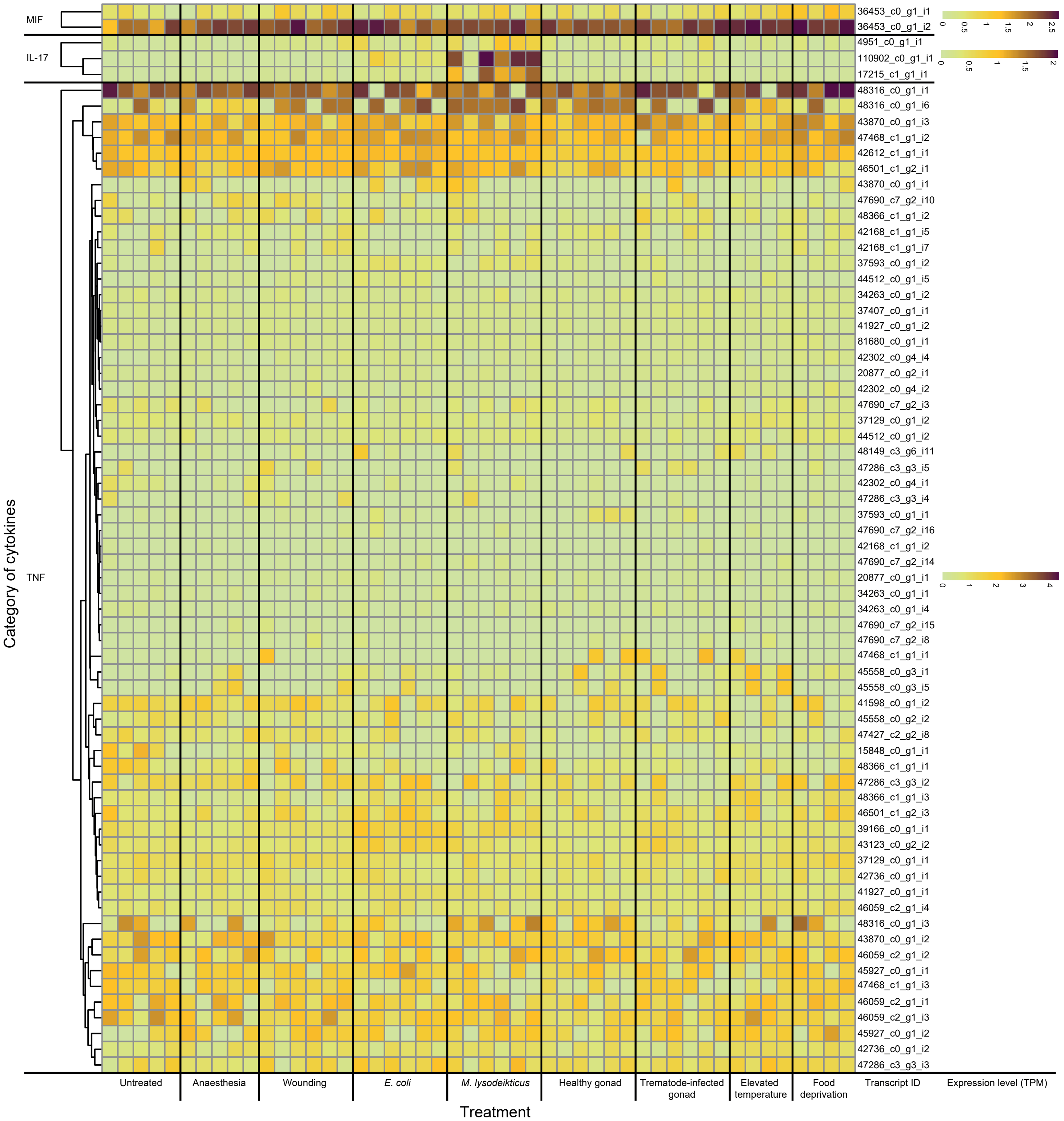

Fig. S6. Expression levels of individual transcripts found to represent annotated cytokines in units of transcripts per million (TPM) for each experimental snail. Heatmap shows the variation for each factor using the dynamic range. Transcripts related to each factor are clustered according to their similarity.

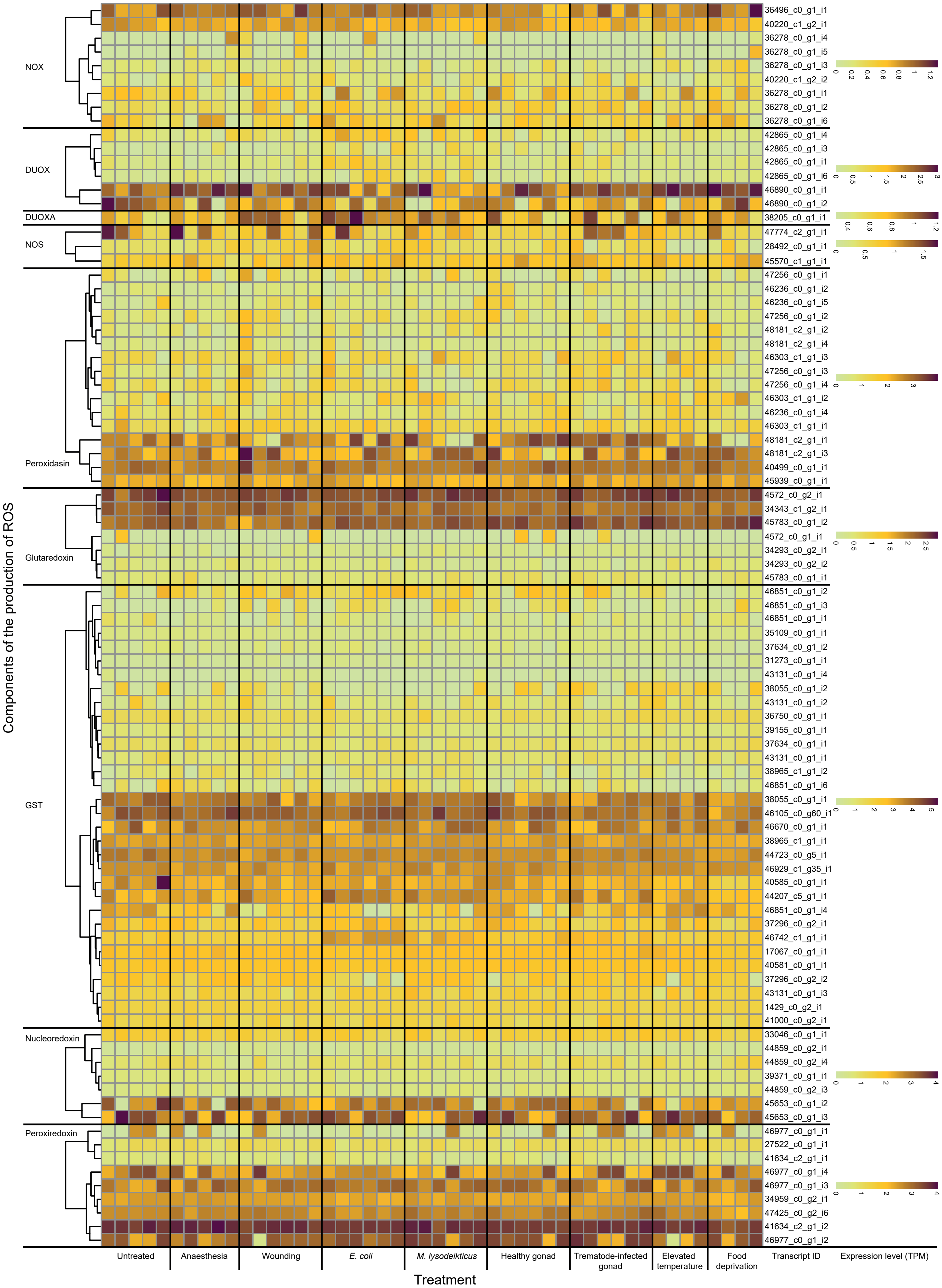

Fig. S7. Expression levels of individual transcripts found to represent annotated factors related to the production of reactive oxygen species (ROS) in units of transcripts per million (TPM) for each experimental snail. Heatmap shows the variation for each factor using the dynamic range. Transcripts related to each factor are clustered according to their similarity.

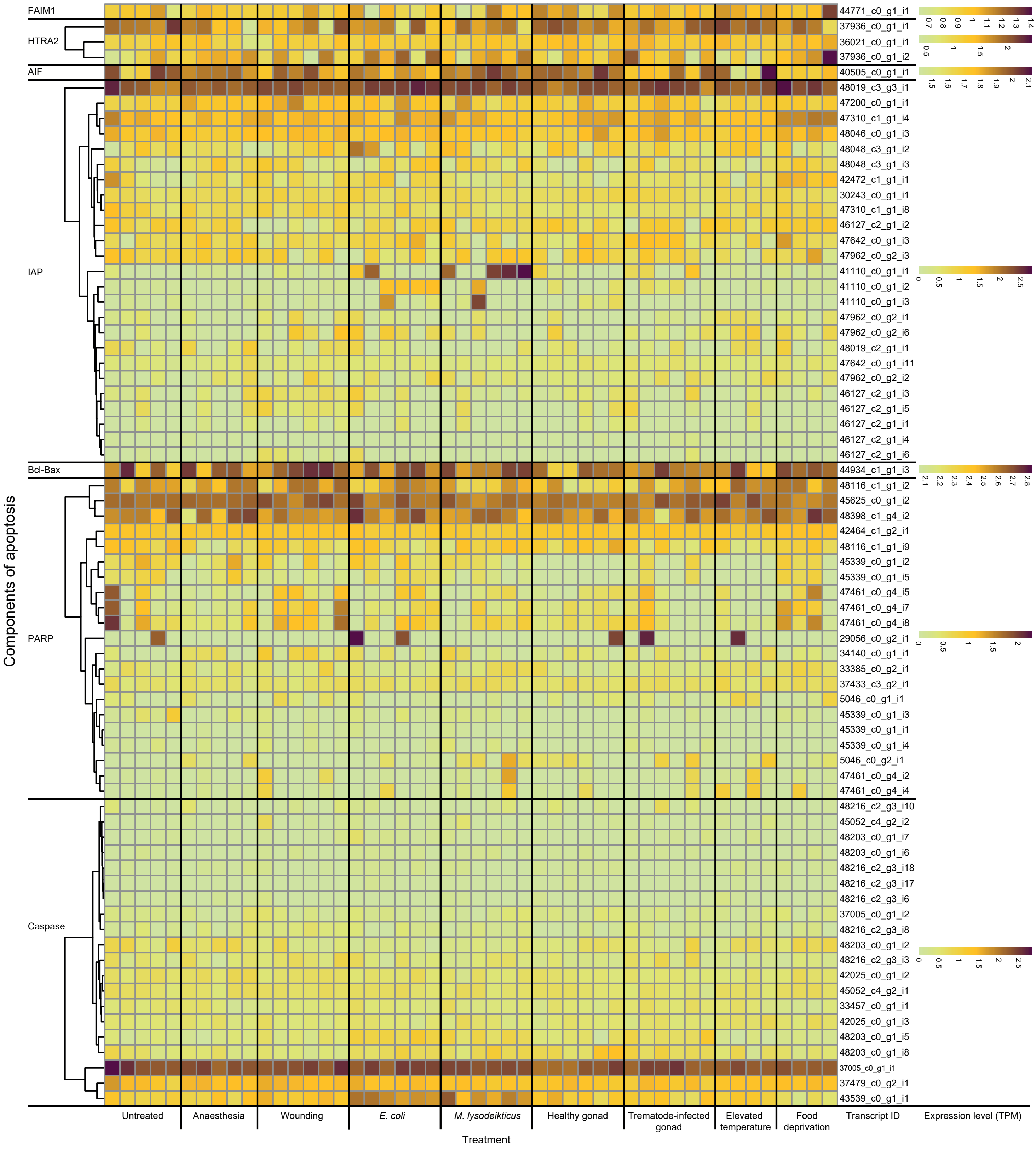

Fig. S8. Expression levels of individual transcripts found to represent annotated factors related to apoptosis in units of transcripts per million (TPM) for each experimental snail. Heatmap shows the variation for each factor using the dynamic range. Transcripts related to each factor are clustered according to their similarity.

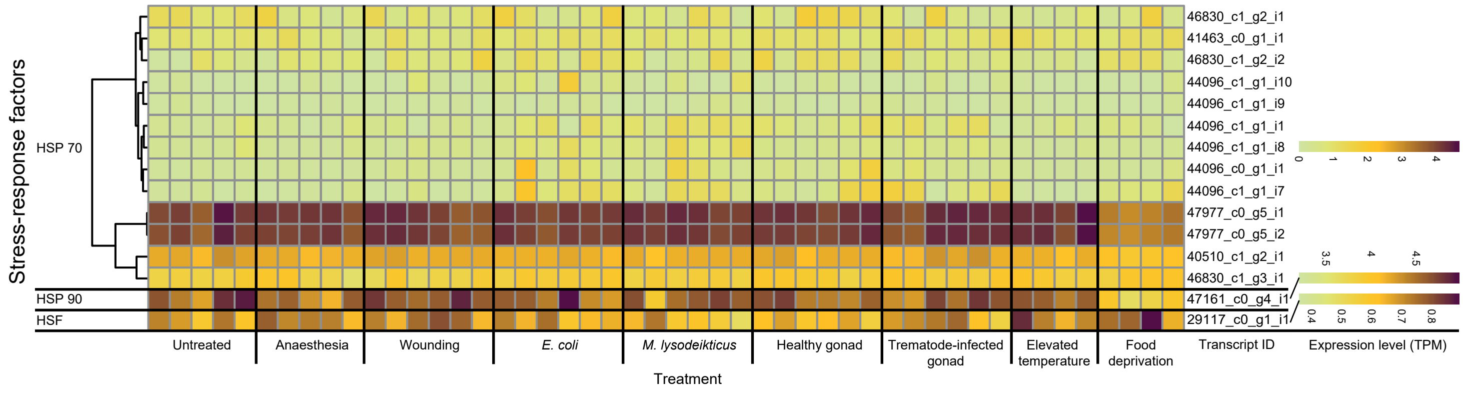

Fig. S9. Expression levels of individual transcripts found to represent the annotated stress-response factors in units of transcripts per million (TPM) for each experimental snail. Heatmap shows the variation for each factor using the dynamic range. Transcripts related to each factor are clustered according to their similarity.

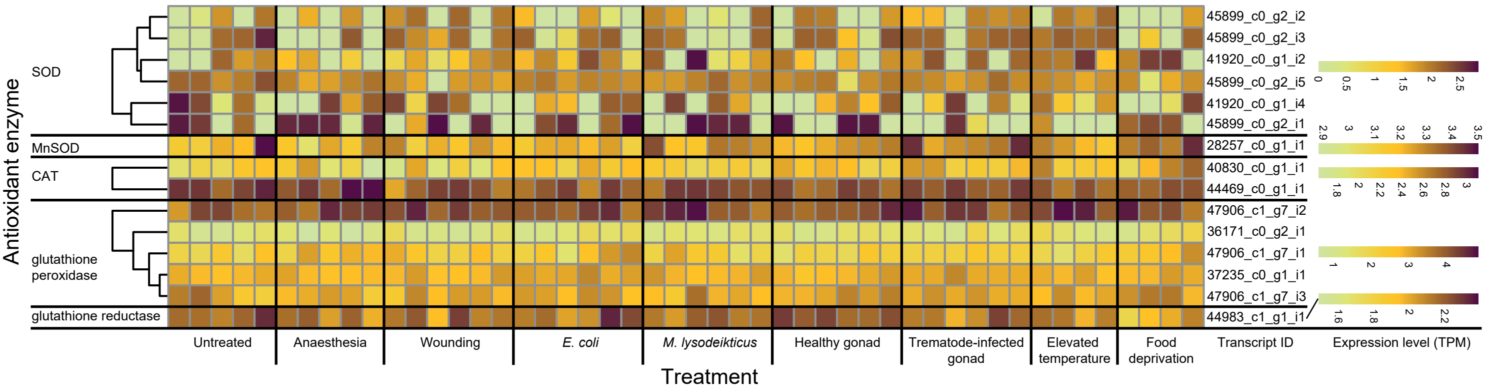

Fig. S10. Expression levels of individual transcripts found to represent the annotated antioxidant enzymes in units of transcripts per million (TPM) for each experimental snail. Heatmap shows the variation for each factor using the dynamic range. Transcripts related to each factor are clustered according to their similarity.

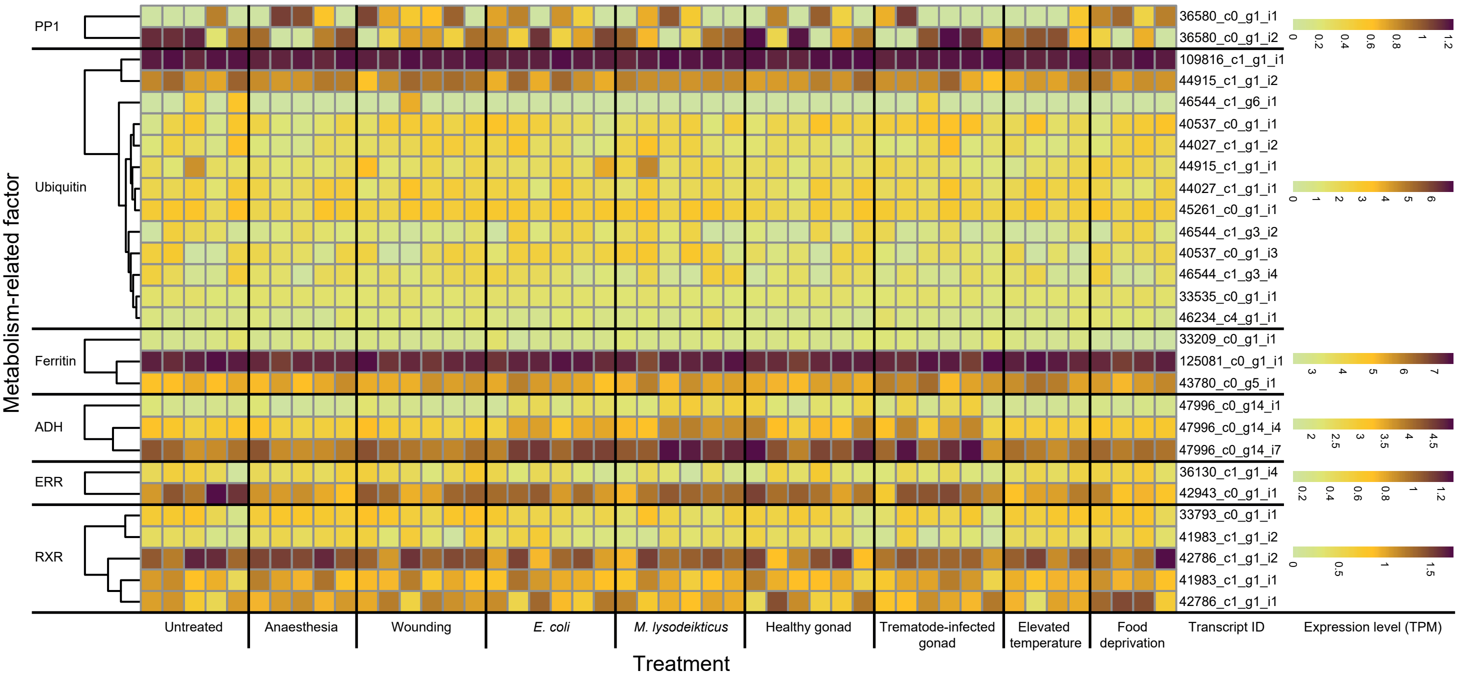

Fig. S11. Expression levels of individual transcripts found to represent annotated factors related to metabolism in units of transcripts per million (TPM) for each experimental snail. Heatmap shows the variation for each factor using the dynamic range. Transcripts related to each factor are clustered according to their similarity.
